# Universal single-copy genes and 16S rDNA present incongruent evolutionary histories in *Vibrio*

**DOI:** 10.64898/2025.12.07.692777

**Authors:** Alexandra García-Flórez, Amaia Leunda-Esnaola, Pablo Arrufat, Vladimir R. Kaberdin, Peter B. Pearman

**Author notes:** **Corresponding author:** V. R. Kaberdin. Department of Biology Education, Svante Arrhenius Väg 20C, Stockholm University, SE-106 91, Stockholm, Sweden. Other author e-mail addresses: Alexandra García-Flórez, Amaia Leunda-Esnaola, Pablo Arrufat, Peter B. Pearman.

## Abstract

A common technique for the study of the diversity and evolution of microbial communities is 16S rDNA sequencing. However, high sequence identity and variable copy number constrain the application of 16S rDNA in differentiation of closely related taxa and estimation of species relative abundance in environmental samples. A promising alternative is the use of universal single-copy genes (USCGs) as phylogenetic markers. We develop this by analyzing a set of USCG loci from the genus *Vibrio*, which holds over 100 species of substantial ecological and epidemiological relevance. The phylogenetic histories of these loci, of representative copies of 16S and 23S rDNA genes, and of a collection of 16S rDNA partial sequences were reconstructed using Bayesian inference. Taxon resolution was assessed according to consensus tree topology and clade credibility values. In addition, the congruence among posterior distributions of phylogenetic estimates of the different loci was calculated using Robinson-Foulds distances and visualized with non-metric multidimensional scaling (NMDS). Phylogenetic analyses reveal that USCG loci produce highly resolved trees in comparison to those of 16S and 23S rDNA sequences. We also observe relatively high congruence among phylogenies of USCG loci while rDNA phylogenies diverge from these. The loci *mfd* and *uvrC* are highlighted for further research on *Vibrio* evolution and analysis of environmental samples. Moreover, possible sources of phylogenetic incongruence between USCG and rDNA loci include differential susceptibility to horizontal gene transfer, as potentially explained by the complexity hypothesis, or lack of phylogenetic information due to limited sequence variability in rDNA sequences.

## INTRODUCTION

Environmental microbiologists have long been challenged to accurately characterize the structure and evolutionary relationships of bacterial communities (Parks et al., 2017). The analysis of microbial communities from environmental samples is often performed with 16S rDNA amplicon-based sequencing (López-Aladid et al., 2023; Větrovský and Baldrian, 2013). However, elevated inter-species similarity at this locus can limit its utility for distinguishing closely related taxa (Bartoš et al., 2024), while substantially divergent copies from the same species can be misclassified (Leunda-Esnaola et al., 2024). Furthermore, horizontal transfer of 16S rDNA segments has been identified, especially at the intra-genus level (Kitahara and Miyazaki, 2013; Schouls et al., 2003; Tian et al., 2015), complicating its use in studies of bacterial communities. The resultingly low phylogenetic resolution can be improved through the use of 23S rDNA, another cistron from the rDNA operon that harbors a greater number of informative bases (Leunda-Esnaola et al., 2024). Nonetheless, the unpredictable number of rDNA operons in each strain can introduce biases into estimates of relative abundances of species in community samples (Starke et al., 2021), while intra-genomic variability among rDNA may obscure the evolutionary structure of bacterial communities (Větrovský and Baldrian, 2013).

To overcome the limitations of 16S rDNA-based analysis, the use of universal single-copy genes (USCGs) has been proposed (Tian and Imanian, 2023; Wang et al., 2022). These genes normally represent protein-coding sequences that are found in most bacterial species. Moreover, these loci generally evolve faster than rDNA, making them potentially more suitable for distinguishing closely related species than is 16S rDNA (Wang et al., 2022; Wu et al., 2013). In addition, some USCGs present low rates of horizontal gene transfer (HGT), suggesting that phylogenetic signal may be conserved (Jain et al., 1999; Parks et al., 2017). Given the promising characteristics of some USCGs as phylogenetic markers, efforts have been also made to determine whether they present congruent phylogenies. Several studies have compared the evolutionary history of 16S rDNA with that of individual USCGs within specific genera (Case et al., 2007; Liu et al., 2021; Maroniche et al., 2017; Wang et al., 2007) primarily focusing on loci that code for the key components of the prokaryotic ribosome and RNA polymerase (Hassler et al., 2022; Wang and Huang, 2018). A more comprehensive analysis examined 148 core genes that have previously been used for phylogenomic analysis and identified the set of 20 core genes that produced the most concordant phylogenies to 16S rDNA evolutionary history in bacteria (Tian and Imanian, 2023).

Despite the successful use of 16S rDNA and some USCGs in phylogenomic analysis, the occurrence and copy number of 16S rDNA, as well as some genes that are frequently considered as USCGs, varies greatly across the bacterial kingdom, making it difficult to resolve species in some groups of prokaryotes, including the *Vibrio* genus (Leunda-Esnaola et al., 2024). The latter comprises important animal and human pathogens, whose occurrence in marine and terrestrial ecosystems can greatly affect animal and human health (Baker-Austin et al., 2018). For instance, the presence of *V. coralliilyticus* can lead to coral bleaching (Kimes et al., 2012), whereas other *Vibrio* pathogens (e.g., *V. alginolyticus* or *V. harveyi*) can cause disease outbreaks in aquaculture settings (Montánchez and Kaberdin, 2020; Sanches-Fernandes et al., 2022). Another notable example is *V. cholerae*, a causative agent of the disease cholera, which remains a public health concern (Bekele et al., 2025). Due to their ecological significance and epidemiological impact, the study of the diversity and evolution of *Vibrio* species is of critical importance (Jesser and Noble, 2018). However, recent studies reveal a number of problems that make detection and classification of environmental *Vibrio* species particularly challenging. In addition to the documented inability of 16S rDNA gene-based trees to reliably identify and delimitate these species (Leunda-Esnaola et al., 2024), other challenges are posed by the high plasticity of *Vibrio* genomes, suggesting that *Vibrio* species are highly recombinogenic and prone to horizontal gene transfer (Torrance et al., 2024). The latter implies that *Vibrio* evolution may largely be dependent on horizontal gene transfer, meaning that even “single-copy” *Vibrio* marker genes are sometimes duplicated, transferred or replaced by xenologs (Eme and Doolittle, 2016; Hehemann et al., 2016). These observations question the capacity of a limited number of USCGs to satisfactory differentiate and delimit *Vibrio* species. Moreover, they highlight the need for examination of the individual evolutionary history of a broader selection of diverse, functionally different USCGs in order to better understand the degree of incongruence between the evolution of these loci and currently used phylogenetic markers.

In this work, we adopt a bioinformatic and evolutionary approach and revisit pre-selected USCG loci (Wang et al., 2022) to assess their individual suitability for phylogenetic analysis of *Vibri*o species. To do this, we apply a Bayesian framework to propose a posterior distribution of evolutionary hypotheses for each locus that is universally present in *Vibrio* genomes. A random sample from each posterior distribution is used to extend the analysis by constructing replicate matrices of the distances among the phylogenies of both USCG and rDNA loci, thus maintaining the variability present in the posterior distributions. We then display the dominant tendencies by constructing two-dimensional ordination diagrams that incorporate both within- and among-locus variability in estimated evolutionary history. This analysis reveals incongruence between evolutionary history of rDNA and USCGs in *Vibrio*. This finding reinforces increasing evidence that the use of 16S rDNA sequences can lead to phylogenetic hypotheses that diverge from those derived from USCGs, which are generally consistent among themselves across the genus *Vibrio*.

## MATERIALS AND METHODS

### Genomes and loci under study

Over 100 different *Vibrio* species have been included into the List of Prokaryotic names with Standing in Nomenclature (LPSN; Parte et al., 2020). On September 12, 2024, we created a database with 87 representative *Vibrio* species that were validly published according to the LPSN database and for which complete chromosome sequences were available on NCBI databases (Supplementary Table S1). In addition, *Escherichia coli* O157:H7 str. Sakai, *Salmonella bongori* NCTC 12419 and *Photobacterium damselae* subsp. *damselae* WMD-P2 were selected as outgroup species for phylogenetic analysis (Supplementary Table S1). Full chromosome sequences and annotations from *Vibrio* species and outgroup taxa were downloaded from NCBI Nucleotide database on September 12, 2024, and March 3, 2025, respectively.

Gene names of USCGs were extracted from the sets of clusters of orthologous genes (COGs) reviewed by Wang et al. (2022) according to the information present in the NCBI COGs Database (Galperin et al., 2021) on September 12, 2024. We identified 125 structurally annotated USCGs in the 87 representative *Vibrio* genomes (data not presented), in addition to various synteny groups composed almost exclusively of USCGs. Full-length sequences of USCGs were retrieved from chromosome records based on existing structural annotations. In case of synteny groups, full-length sequences were extracted with SAMtools v1.21 (Danecek et al., 2021) using their terminal coordinates in each genome. The 25 longest USCGs (> 1500 bp) and the synteny groups were evaluated for Bayesian phylogenetic reconstruction (see below), and only the loci that met our diagnostic criteria were selected for this study. These loci included 16 long USCGs, five synteny groups and a synteny subgroup (Supplementary Table S2). Discarded loci are reported in Supplementary Table S3. All USCGs under study were classified according to functional annotations from the NCBI COG Database on September 12, 2024.

### Retrieval and selection of 16S and 23S rDNA sequences

All annotated 16S and 23S rDNA sequences from the 87 *Vibrio* species and the three outgroup taxa were retrieved from chromosome records using a bash script and were classified into pairs according to rDNA operon membership. The 16S rDNA copies of each species were aligned using MAFFT v7.526 (Katoh and Standley, 2013) following a global alignment strategy (G-INS-i) to create 90 multiple sequence alignments (87 *Vibrio* species and three outgroup species). Matrices of pairwise distances between the sequences within each of the 90 alignments, calculated as the proportion of variable sites, were computed using the *dist.dna* function from the *ape* R package v5.8-1 (Paradis and Schliep, 2019). Distance matrices were projected into 2-dimensional space with non-metric multidimensional scaling (NMDS; Kruskal, 1964) using the *monoMDS* function from the *vegan* R package v2.6-8 (Oksanen et al., 2025), with *weakties* set to *False*. The same procedure was followed for the 23S rDNA sequences of each species. Based on the NMDS results, operons with outlier 16S and/or 23S rDNA copies were discarded. Then, for each species, a single rDNA operon was randomly selected and its 16S and 23S rDNA sequences were used as representative copies (Supplementary Table S1). We were also interested in which regions of 16S rDNA might be responsible for any variability in phylogenetic topologies. To address this, we employed universal PCR primer sets (Supplementary Table S4) to delimit variable regions (V), and retrieved 15 partial 16S rDNA sequences from the 16S rDNA representative copies, each of which lacked one or more variable regions (Supplementary Table S5).

### Alignment of *Vibrio* and outgroup sequences

Sequences of the rDNA and USCG loci from the 87 *Vibrio* species and three outgroup taxa were aligned with MAFFT using the G-INS-i algorithm. The number of variable bases of each multiple sequence alignment was calculated with MEGA 11 v11.0.13 (Tamura et al., 2021).

### Bayesian phylogenetic reconstruction

MrBayes v3.2.7 (Ronquist et al., 2012) was used to perform Bayesian phylogenetic inference with the Metropolis-coupled Markov Chain Monte Carlo (MC^3^) algorithm (Geyer, 1991; Yang and Rannala, 1997). To evaluate substitution models, Modeltest-NG v0.1.7 (Darriba et al., 2020) was run with the flags *-d nt -t ml -T mrbayes*, and the best-fit model according to the Bayesian information criterion (BIC) was chosen for each alignment (Supplementary Table S6).

Inference was conducted with a relaxed molecular clock and the independent gamma rates (IGR) model in MrBayes v3.2.7 (Lepage et al., 2007). A gamma distribution prior with shape parameter α = 1 and rate parameter λ = 1 was chosen for the basal clock rate, expressed as expected substitutions per site per million years (Duchêne et al., 2016; Gibson and Eyre-Walker, 2019). Enterobacteria were set as a sister to Vibrionaceae, and *Photobacterium damselae* was set as a sister to *Vibrio* (Lin et al., 2018). We used a gamma distribution with mean and standard deviation measured in millions of years ago (MYA) for time calibration of internal nodes. Node dating was configured as follows: 796 ± 123.72 MYA for the root node (Battistuzzi et al., 2004), 568 ± 32 MYA for the divergence between *Photobacterium* and *Vibrio* (Lin et al., 2018; Sawabe et al., 2007), and 130 ± 30 MYA for the divergence between *Escherichia* and *Salmonella* (Doolittle et al., 1996; Ochman and Wilson, 1987). All tree tips were set to exist in the present.

For each alignment, we used four independent MrBayes runs with five chains each, and set burn-in to 40%. Run parameters were configured to obtain adequate posterior distributions according to the following criteria: (1) the performance of the MC^3^ algorithm was acceptable when chain swap frequencies between adjacent chains were between 10% and 70% (Ronquist et al., 2020); (2) convergence was acceptable when the average standard deviation of split frequencies (ASDSF) (Lakner et al., 2008) was below 0.03 (Ronquist et al., 2020) and when the average potential scale reduction factor (PSRF) was below 1.1 (Gelman et al., 2013; Gelman and Rubin, 1992); and finally, (3) sampling was acceptably efficient when the effective sample size (ESS) for all parameters exceeded 200 (Rambaut et al., 2018), and when no elevated autocorrelation existed in the trace of the logarithm of the posterior likelihood (LnL), which we visualized in Tracer v1.7.2 (Rambaut et al., 2018). Particularly, the temperature constant was lowered from 0.1 to 0.075, 0.05 or 0.025 to meet criterion (1), and the number of generations and sample frequency were extended from 5 x 10e7 and 2 x 10e4, respectively, to 1 x 10e8 and 4 x 10e4, respectively, if criteria (2) and (3) were not met. Despite extensive adjustment of parameters, 22 USCG alignments did not produce stable, convergent results during MC^3^ runs, hence they were excluded from this study (Supplementary Table S3). We report results using posterior tree distributions from all alignments with convergent results (for final configurations see Supplementary Table S6). The results of phylogenetic inference were summarized in a 50% majority rule consensus tree with clade credibility values expressed as posterior branch probabilities. We considered bipartitions as completely supported when their credibility values were 1.

### Distances between phylogenetic trees

We randomly selected a total of 999 phylogenetic trees from each posterior distribution produced from the rDNA and USCG alignments. Branch lengths were converted to time units (million years) using the *obtainDatedPosteriorTreesMrB* function from the *paleotree* R package v3.4.7 (Bapst, 2012), and trees were rescaled to a tree length of 1 (Matzke et al., 2014). The normalized Robinson-Foulds (nRF) distance (Day, 1985; Robinson and Foulds, 1981) was used to quantify topological dissimilarity among trees produced from the alignments, and the weighted Robinson-Foulds (wRF) distance (Robinson and Foulds, 1979) was used to include relative differences in node depth into the evaluation of tree dissimilarity. In both cases, we generated 999 pairwise distance matrices by choosing without replacement from the 999 trees sampled from the posterior distribution that was produced from each alignment. The nRF and wRF distances were computed employing the *RF.dist* and *wRF.dist* functions from the *phangorn* R package v2.12.1 (Schliep, 2011).

### Analysis of phylogenetic congruence through NMDS

The matrices of nRF and wRF distances were transformed into 2-dimensional representations using the *monoMDS* function from the *vegan* R package and aligned with general Procrustes analysis (Commandeur, 1991) as implemented in the *FactoMineR* R package v2.11 (Lê et al., 2008). The mean position and standard deviation of each alignment in both dimensions were calculated from the set of 999 configurations after Procrustean transformation. This made the position of all trees comparable in the space of the two NMDS axes.

## RESULTS

### Description of USCG loci

The focal loci chosen for this study (Supplementary Table S2) include 36 USCGs (Supplementary Table S7), 18 of which are related to translation, ribosomal structure and biogenesis (J). Nine USCGs code for ribosomal proteins and four USCGs code for aminoacyl-tRNA synthetases. Other functional categories are represented, including replication, recombination and repair (L), and transcription (K). Most synteny groups harbor genes related to the same function, such as replication, recombination and repair in case of group D, or ribosomal structure in case of group E. The USCG loci are located on chromosome I in 84 *Vibrio* species, but on chromosome II in *V. alginolyticus*, *V. fluvialis*, *V. rotiferianus* and *V. zhugei*. The only exceptions are *aspS* and group D in *V. astriarenae*, *polA* in *V. aestuarianus*, and *nrdA* in *V. cincinnatiensis*. Another notable characteristic is that the number of variable bases was higher in the case of the USCG loci than in the rDNA sequences (Supplementary Table S8).

### Phylogenetic inference and congruence

For each alignment, convergence and efficient sampling are evaluated according to various diagnostic metrics (Supplementary Table S9). In most cases, consensus phylogenies are not fully resolved and present one or two trichotomies (Table 1). Furthermore, not all informative bipartitions (i.e., clades composed of more than one taxon) are completely supported (Table 1). Overall, the worst performing alignments are 16S rDNA and its partial sequences, which result in consensus trees with various multifurcations and few completely supported informative bipartitions.

**Table 1.** Characteristics of the consensus trees obtained through Bayesian inference.

| Alignment | Topological characteristics | Number of completely supported informative bipartitions |
| --- | --- | --- |
| 16S rDNA | 9 polytomies with $\geq 3$ descendants | 24 |
| 16S rDNA V1-V2+V4-V9 | 12 polytomies with $\geq 3$ descendants | 19 |
| 16S rDNA V1-V2+V7-V9 | 9 polytomies with $\geq 3$ descendants | 12 |
| 16S rDNA V1-V3+V5-V9 | 11 polytomies with $\geq 3$ descendants | 17 |
| 16S rDNA V1-V5 | 13 polytomies with $\geq 3$ descendants | 17 |
| 16S rDNA V1-V6 | 12 polytomies with $\geq 3$ descendants | 20 |
| 16S rDNA V1-V6+V8-V9 | 11 polytomies with $\geq 3$ descendants | 23 |
| 16S rDNA V1-V7 | 6 polytomies with $\geq 3$ descendants | 22 |
| 16S rDNA V1-V7+V9 | 8 polytomies with $\geq 3$ descendants | 20 |
| 16S rDNA V1-V8 | 11 polytomies with $\geq 3$ descendants | 25 |
| 16S rDNA V2-V9 | 9 polytomies with $\geq 3$ descendants | 18 |
| 16S rDNA V3-V4 | 13 polytomies with $\geq 3$ descendants | 6 |
| 16S rDNA V3-V6 | 13 polytomies with $\geq 3$ descendants | 12 |
| 16S rDNA V3-V7 | 13 polytomies with $\geq 3$ descendants | 12 |
| 16S rDNA V3-V9 | 13 polytomies with $\geq 3$ descendants | 15 |
| 16S rDNA V4-V9 | 13 polytomies with $\geq 3$ descendants | 13 |
| 23S rDNA | 7 trichotomies | 47 |
| *alaS* | Fully resolved | 65 |
| *aspS* | 3 trichotomies | 62 |
| *glnS* | 5 polytomies with $\geq 3$ descendants | 49 |
| *gyrB* | 4 polytomies with $\geq 3$ descendants | 59 |
| *leuS* | 1 trichotomy | 51 |
| *metG* | 3 polytomies with $\geq 3$ descendants | 49 |
| *mfd* | 1 trichotomy | 69 |
| *nrdA* | 2 trichotomies | 56 |
| *parC* | Fully resolved | 61 |
| *parE* | 1 trichotomy | 54 |
| *pnp* | 2 trichotomies | 61 |
| *polA* | Fully resolved | 63 |
| *recN* | 1 trichotomy | 55 |
| *rpoB* | 2 polytomies with $\geq 3$ descendants | 62 |
| *typA* | 3 polytomies with $\geq 3$ descendants | 53 |
| *uvrC* | 2 trichotomies | 53 |
| Group A | 1 trichotomy | 72 |
| Group A.1 | 1 trichotomy | 52 |
| Group B | Fully resolved | 60 |
| Group C | 2 trichotomies | 55 |
| Group D | 1 trichotomy | 51 |
| Group E | 2 trichotomies | 51 |

In the NMDS analysis of tree distances to assess phylogenetic congruence among rDNA and the USCG loci (Fig. 1; Supplementary Table S10), most USCG loci cluster together, with the exception of the outlier locus *typA* (Fig. 1A). Furthermore, trees with originally low total tree length (Supplementary Fig. S1), such as those produced from the alignments of groups A and E, appear in a central position within the cluster when phylogenetic incongruence is quantified according to the nRF distance (Fig. 1A), but on the ordination periphery when branch lengths are additionally considered according to the wRF distance (Fig. 1B). Some genes, for example *mfd* and *uvrC,* appear in the main cluster in both nRF and wRF ordinations (Fig. 1). In addition, 16S rDNA appears as an outlier in comparisons using both distance measures (Fig. 1). When considering only tree topology, trees produced from 23S rDNA alignment are more similar to those generated by USCG loci than are trees deriving from 16S rDNA (Fig. 1A). Nonetheless, trees representing 23S rDNA appear as an outlier when branch lengths are additionally considered (Fig. 1B). Ordination of additional trees representing the selected 16S rDNA copies with single variable regions removed (Fig. 2; Supplementary Table S11) reveals that these variants cluster with the phylogenetic trees produced by the full-length 16S rDNA alignment. Hence, deletion of a single variable region does not here markedly affect the phylogenetic congruence of the 16S rDNA gene with USCG loci.

**Fig. 1.**
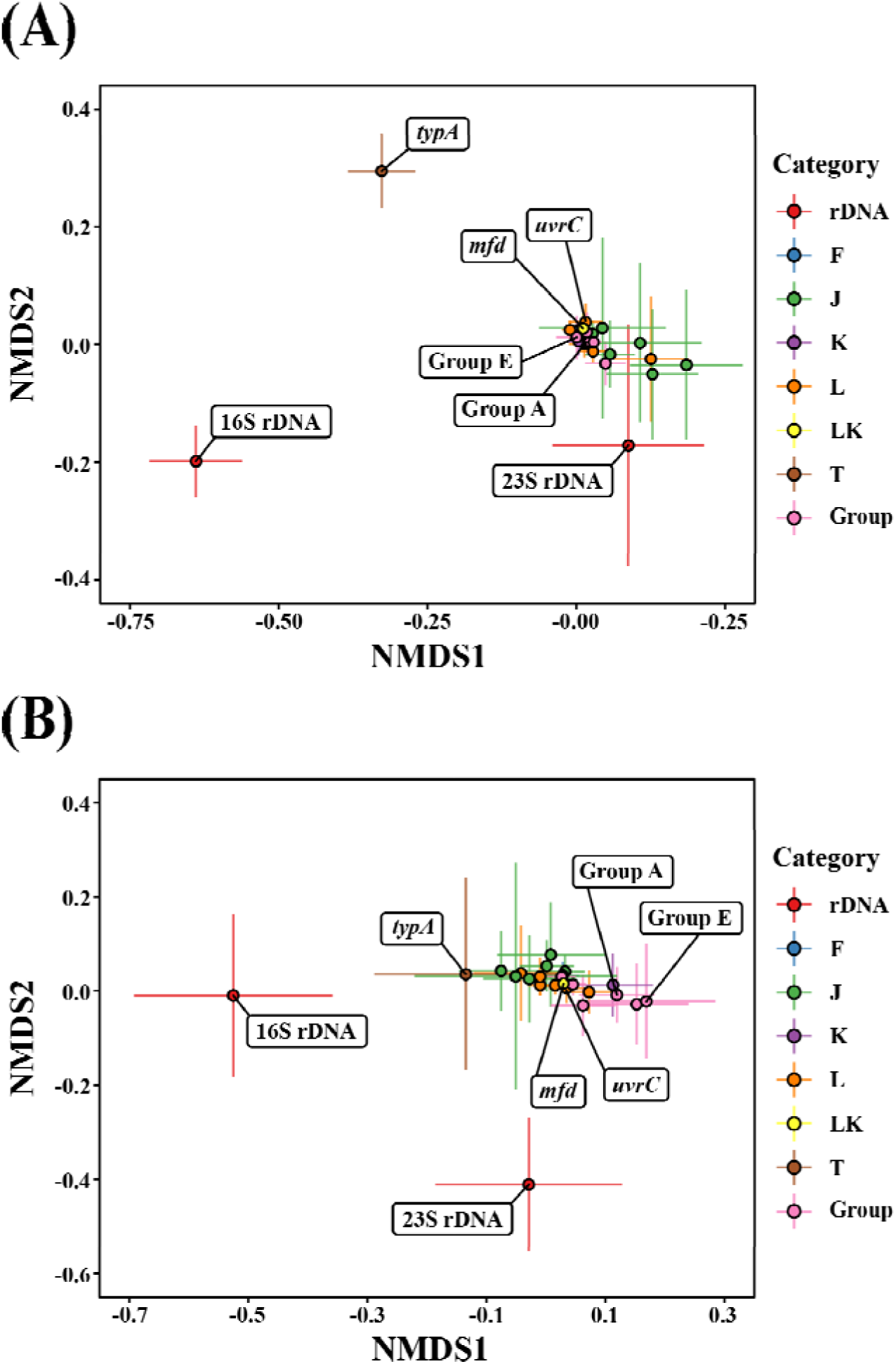
Non-metric multidimensional scaling (NMDS) analysis of phylogenetic congruence between USCG loci and rDNA representative copies. Mean position of loci (dots) with standard deviation bars in two dimensions. **(A)** Ordination of distances among trees calculated according to the nRF metric, which reflects variation in tree topologies. Stress (mean ± standard deviation): 0.13 ± 0.01. **(B)** Ordination of distances among trees calculated according to the wRF metric, which additionally reflects variation in tree branch lengths. Stress (mean ± standard deviation): 0.15 ± 0.02. Categories: rDNA represents 16S and 23S rDNA loci; F, J, K, L, LK and T represent individual USCGs according to their functional categories (Supplementary Table S7); and synteny groups as indicated.

**Fig. 2.**
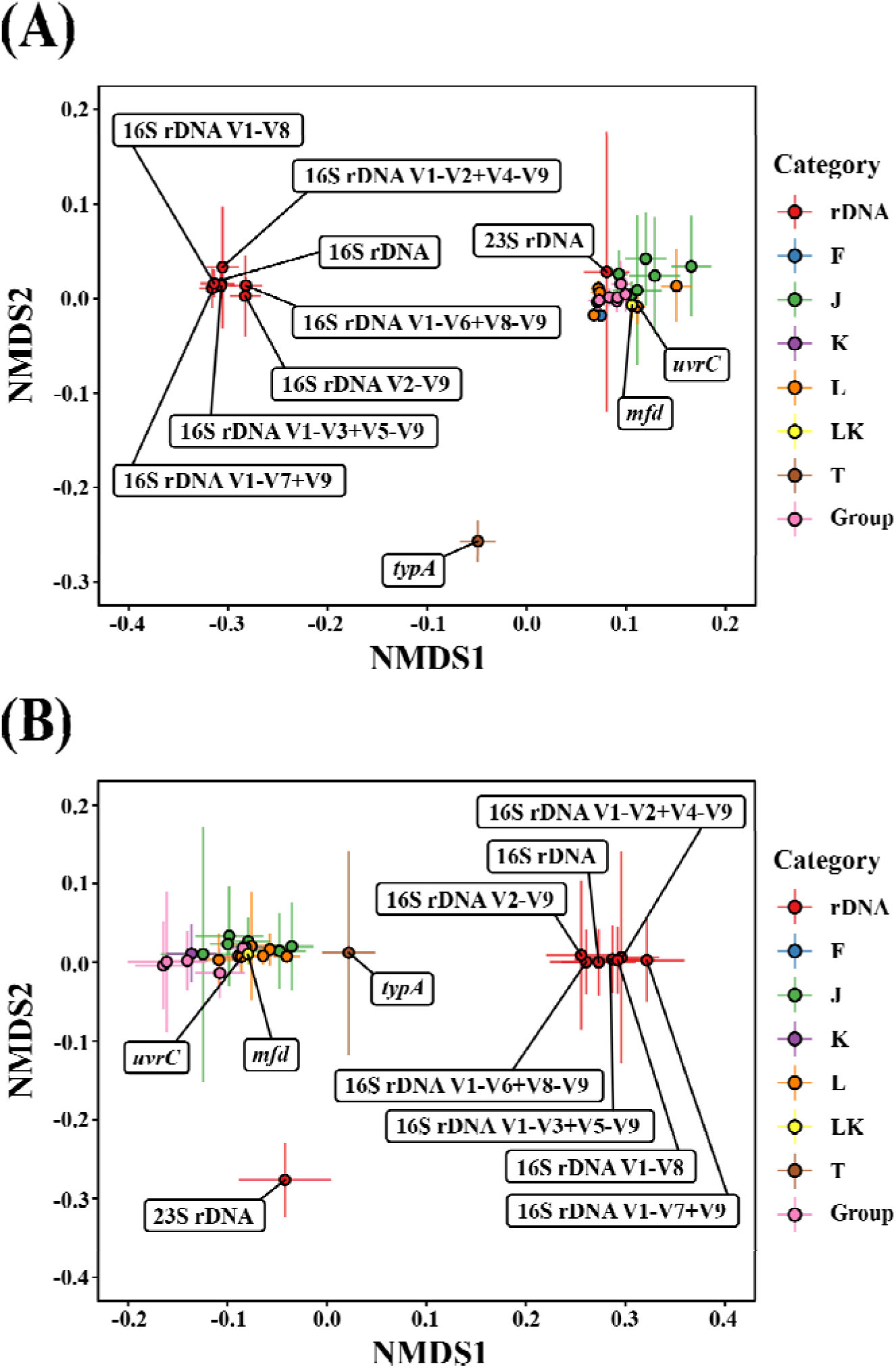
NMDS analysis of phylogenetic congruence between USCG loci, rDNA representative copies, and partial 16S rDNA sequences without a single variable region. Mean position of loci (dots) with standard deviation bars in two dimensions. **(A)** Ordination of distances among trees calculated according to the nRF metric. Stress (mean ± standard deviation): 0.09 ± 0.01. **(B)** Ordination of distances among trees calculated according to the wRF metric. Stress (mean ± standard deviation): 0.12 ± 0.02. Categories: rDNA represents 16S and 23S rDNA loci; F, J, K, L, LK and T represent individual USCGs according to their functional categories (Supplementary Table S7); and synteny groups as indicated.

Finally, we also ordinated phylogenetic trees produced by rDNA representative copies and USCG loci together with a collection of 16S rDNA variants lacking one or more variable regions (Fig. S2; Supplementary Table S12). Partial 16S rDNA sequences form a dispersed cluster together with 16S rDNA representative copies, highlighting that the deletion of multiple variable regions can visibly impact phylogenetic incongruence among loci. Particularly, some variants containing V3-V4, such as 16S rDNA V1-V5, 16S rDNA V3-V4, 16S rDNA V3-V6 and 16S rDNA V3-V7, appear in both ordination plots relatively distant from trees produced by USCG loci, compared to trees of 16S rDNA evolution (Fig. S2).

## DISCUSSION

Recent studies highlight the shortcomings of using 16S rDNA sequences for phylogenetic and environmental studies when working at the intra-genus level (Dewhirst et al., 2005; Liu et al., 2021; Vos et al., 2012; Yang et al., 2022). Given these limitations, the use of USCG loci provides an alternative that can potentially improve phylogenetic resolution and species differentiation (Liu et al., 2021; Vos et al., 2012; Yang et al., 2022). In the present study, 22 *Vibrio* single-copy loci (Supplementary Table S2) show promising results in comparison to 16S rDNA for their use in phylogenetic studies.

Although these protein-coding loci resolve phylogenetic relationships with variable success, their performance is notably higher than that of 16S rDNA (Table 1). This has been observed in another study, where the phylogenetic resolution provided by 16S rDNA is compared to that of *gyrB* in Myxococcaceae (Liu et al., 2021). Nonetheless, other loci analyzed here yield improvements in clade resolution and node support over the use of either 16S rDNA or *gyrB*, for example *alaS*, *mfd*, *parC*, *polA*, and groups A and B. While 23S rDNA offers better resolution than 16S rDNA, as noted in Leunda-Esnaola et al. (2024), it does not outperform the USCG loci analyzed in the present study. Although multiple substitutions can introduce noise (Simmons et al., 2004), differences in the number of variable sites between rDNA and USCG loci (Supplementary Table S8) probably contribute to differences in phylogenetic resolution.

Phylogenetic congruence among the posterior distributions of trees produced by each alignment is here visualized through procrustean analysis of NMDS ordinations. In topological comparisons of phylogenetic trees (Fig. 1A; Supplementary Table S10), most USCG loci cluster together in ordination space, while 23S rDNA trees appear in the periphery of the cluster and 16S rDNA shows elevated phylogenetic incongruence with the other loci. Topological incongruence can reflect differential rates of HGT (Steenwyk et al., 2023). Matzke et al. (2014) and Lerat et al. (2003) report high phylogenetic congruency and little HGT in a set of marine cyanobacteria and Gammaproteobacteria core proteins, respectively. Our results suggest high phylogenetic consistency among the USCG loci analyzed in the present study in the genus *Vibrio* and potentially imply low rates of HGT in these genes (Jain et al., 1999; Parks et al., 2017). This conclusion could be supported by the complexity hypothesis, which states that genes involved in information processes (i.e., replication, transcription and translation) are less susceptible to HGT due to the involvement of their protein products in large complexes (Jain et al., 1999; Rivera et al., 1998). A recent study further supports this idea (Martinez-Gutierrez and Aylward, 2021).

The relative position of phylogenetic trees in the ordination space constructed with wRF index dissimilarity (Fig. 1B; Supplementary Table S10) depends on differences in internal node depths. Alignments whose posterior distribution of trees originally had the lowest total tree length (Supplementary Fig. S1) tend to appear at the periphery of the cluster of USCG points (Fig. 1B; Supplementary Table S10). Matzke et al. (2014) also report this phenomenon when rescaling non-ultrametric trees with branch lengths measured in number of substitutions per site, and they suggest the amount of information present in short branches as a possible cause. Hence, our results help shed light on the behavior of this metric. Overall, alignments that show centrality in both ordinations (Fig. 1) could be concatenated in order to produce highly supported trees that could reflect the evolutionary history of a larger part of the *Vibrio* genome (Martinez-Gutierrez and Aylward, 2021).

The use of the 16S rDNA alignment produces phylogenetic trees that differ from the evolutionary histories of the chosen USCG loci (Fig. 1; Supplementary Table S10). Horizontal gene transfer of 16S rDNA has been previously observed between closely related taxa (Schouls et al., 2003; Tian et al., 2015). These transfers most likely involve sequence segments rather than full-length copies, thus diluting phylogenetic signal without compromising ribosomal structure and function (Sato and Miyazaki, 2017; Wang and Zhang, 2000). To address this, we report ordinated phylogenetic trees from 16S rDNA representative sequences that lack one variable region (Fig. 2; Supplementary Table S11), but the results were not completely conclusive. Potentially, factors leading to phylogenetic incongruence within this genus could span a larger region of the 16S rDNA gene, or even be present throughout its sequence. To gain further insight into this issue, we analyze trees from a variety of 16S rDNA alignments that lacked multiple variable regions (Supplementary Fig. S2; Supplementary Table S12). This reveals higher variability among their phylogenetic histories than that seen as a result of single-region deletions. Interestingly, phylogenetic incongruence among different 16S rDNA segments has also been reported. For instance, in *Escherichia* and *Shigella*, 16S rDNA V1-V2, V1-V3, V4 and V3-V5 partial sequences show varying phylogenetic congruence with the evolutionary history of 16S rDNA. Among these segments, the V1-V3 region is the most concordant with the phylogenetic tree produced by the full-length 16S rDNA sequence (Johnson et al., 2019). In our data, alignments containing the 16S rDNA V3-V4 region produce phylogenetic trees that greatly diverge from USCG evolutionary history in *Vibrio* (Supplementary Fig. S2; Supplementary Table S12). This has been observed in other genera. For example, Hrovat et al. (2024) analyze a wide range of plant-associated bacteria, including *Enterobacter* and other Gammaproteobacteria, and conclude that phylogenies based on USCGs and on average nucleotide identity (ANI) are congruent. In contrast, phylogenies based on 16S rDNA V3-V4 and V4 regions are not as similar to ANI phylogeny as are those phylogenies produced from other 16S rDNA segments.

The 16S rDNA V3-V4 region could be further studied to detect potential accumulation of HGT or other sources of incongruence in *Vibrio* that could explain our results. For instance, González-Escalona et al. (2005) observe that a segment of 16S rDNA sequence within the V3-V4 region harbors intragenomic and intergenomic variability among various strains in *V. parahaemolyticus*, suggesting that HGT affects this segment. Transfers within this region have also been proposed between *V. splendidus* and *Colwellia* (Jensen et al., 2009). Furthermore, in other Gammaproteobacteria, such as *Enterobacter*, 16S rDNA recombination among strains has also been proposed, with a suggested transferred region overlapping with the 16S rDNA V3-V4 segment (Sato and Miyazaki, 2017). Nonetheless, phylogenetic incongruence of full-length and partial 16S rDNA sequences could also be attributed to low phylogenetic resolution (Table 1). The limited variability among rDNA sequences in comparison to USCGs (Supplementary Table S8) could hinder their use to differentiate closely related species, leading to errant hypotheses of evolutionary history due to insufficient phylogenetic information (Hrovat et al., 2024; Johnson et al., 2019)

We observe that 23S rDNA phylogenies diverge from trees produced by 16S rDNA sequences, but fail to represent USCG evolution (Fig. 1; Supplementary Table S10). Similarly, Leunda-Esnaola et al. (2024) highlight that *Vibrio* 16S and 23S rDNA sequences produce topologically dissimilar trees due to differences in the amount of phylogenetic information contained in each locus. Furthermore, Dewhirst et al. (2005) report that phylogenies of 23S rDNA are more congruent with the evolutionary history of 870 proteins than with the phylogeny of 16S rDNA in *Helicobacter*. However, research on phylogenetic incongruence sources involving 23S rDNA, such as HGT, is less extensive than in the case of 16S rDNA.

In addition to phylogenetic considerations, the location of USCGs within *Vibrio* genomes merits a brief examination. Given the essential functions performed by these genes, it is surprising that all of them are annotated on chromosome II in four *Vibrio* species. However, these assemblies representing chromosome II have the typical size and gene set of chromosome I, and vice versa (Kirkup et al., 2010), suggesting that the chromosomes may have been misannotated in these four strains. Further research could resolve this issue.

Ultimately, the findings of this study are largely consistent with the existing literature on bacteria in other families and genera, and support our initial hypotheses concerning the potential utility of USCG loci in phylogenetic analyses. In the present study some genes yield particularly promising results according to high branch support and congruence. For instance, *mfd* and *uvrC* show high phylogenetic resolution (Table 1) and relatively high congruence with most USCG loci (Figs. 1, 2; Supplementary Fig. S2). This suggests that concatenation of various USCGs could produce highly supported trees. The utility of using these genes for biodiversity studies of microbe community composition based on amplicon-sequencing merits further research and experimental validation.

## CONCLUSIONS

In this work, we evaluate various USCG loci in the *Vibrio* genome in terms of phylogenetic resolution and congruence. We find highly supported consensus trees and substantial clustering in ordination space in comparison with full-length and partial 16S rDNA sequences. After a closer examination of the outcomes of these analyses, some genes are proposed for deeper phylogenetic study through alignment concatenation and the development of focal loci for the study of microbe community composition.

## CRediT AUTHORSHIP CONTRIBUTION STATEMENT

**Alexandra García-Flórez**: Data curation, Formal analysis, Software, Validation, Visualization, Writing – Original draft, Writing – Review & Editing. **Amaia Leunda-Esnaola**: Data curation, Writing – Review & Editing. **Pablo Arrufat**: Visualization, Writing – Review & Editing. **Vladimir Kaberdin**: Conceptualization, Funding acquisition, Methodology, Project administration, Supervision, Writing – Review & Editing. **Peter B. Pearman**: Conceptualization, Data curation, Funding acquisition, Methodology, Project administration, Supervision, Writing – Review & Editing.

## DATA STATEMENT

Multisequence alignments for all loci analyzed in this study are available via the following URL: https://doi.org/10.5281/zenodo.17789687

## ACKNOWLEDGEMENTS

We are thankful for the technical support provided by the Scientific Computing Service (IZO-SGI) unit of General Research Services (SGIker) of the University of the Basque Country (UPV/EHU) and European funding (ERDF and ESF). This research was conducted in partial fulfillment of the requirements for conferral of a Bachelor’s Degree by the University of the Basque Country to A.G.-F.

