## Supplemental materials for "Universal single-copy genes and 16S rDNA present incongruent evolutionary histories in *Vibrio*"

**This file includes:**

- Supplementary Tables S1-S12.
- Supplementary Figures S1-S2.
- Supplementary References

**Supplementary Table S1. Representative strains and their cognate 16S and 23S rDNA loci analyzed in this study.** Genome assembly and rDNA loci names are provided according to NCBI nomenclature.

| **Representative Strain** | **Genome Assembly** | **16S rDNA locus** | **23S rDNA locus** |
| --- | --- | --- | --- |
| *Vibrio aerogenes* LMG 19650 | GCF_024346755.1 | OCV29_RS00835 | OCV29_RS00850 |
| *Vibrio aestuarianus* subsp. *francensis* 02/041 | GCF_012395215.1 | HGG77_RS01620 | HGG77_RS01605 |
| *Vibrio alfacsensis* 04Ya108 | GCF_019670485.1 | VA249_RS12810 | VA249_RS12795 |
| *Vibrio algicola* SM1977 | GCF_009601765.2 | GFB47_RS11180 | GFB47_RS11165 |
| *Vibrio alginolyticus* E110 | GCF_023650915.1 | NAL94_RS08775 | NAL94_RS08780 |
| *Vibrio anguillarum* NB10 | GCF_000786425.1 | VANGNB10_RS06640 | VANGNB10_RS06655 |
| *Vibrio aphrogenes* CA-1004 | GCF_002157735.2 | VCA1004_RS00590 | VCA1004_RS00610 |
| *Vibrio artabrorum* CECT 7226 | GCF_024347295.1 | OCU36_RS14805 | OCU36_RS14780 |
| *Vibrio astriarenae* HN897 | GCF_010587385.1 | GT360_RS00685 | GT360_RS00700 |
| *Vibrio atlanticus* LGP32 | GCF_000091465.1 | VS_RS20605 | VS_RS20575 |
| *Vibrio azureus* LC2-005 | GCF_002849855.1 | BS333_RS02620 | BS333_RS02635 |
| *Vibrio breoganii* CAIM 1829 | GCF_024346795.1 | OCU35_RS01400 | OCU35_RS01420 |
| *Vibrio campbellii* BoB-53 | GCF_002906475.1 | C1N50_RS16305 | C1N50_RS16275 |
| *Vibrio casei* DSM 22364 | GCF_002218025.2 | VCASEI_RS12410 | VCASEI_RS12395 |
| *Vibrio celticus* CECT 7224 | GCF_024347335.1 | OCV19_RS00310 | OCV19_RS00315 |
| *Vibrio chagasii* LMG 21353 | GCF_024347355.1 | OCV52_RS15535 | OCV52_RS15510 |
| *Vibrio cholerae* RFB16 | GCF_008369605.1 | F0316_RS01805 | F0316_RS01820 |
| *Vibrio cidicii* 2756-81 | GCF_009763805.1 | GPY24_RS19080 | GPY24_RS19065 |
| *Vibrio cincinnatiensis* 1398-82 | GCF_009763485.1 | GPX87_RS06175 | GPX87_RS06195 |
| *Vibrio comitans* LMG 23416 | GCF_024346835.1 | OCU46_RS00505 | OCU46_RS00520 |
| *Vibrio coralliilyticus* Rb102 | GCF_029541605.1 | P6988_RS00610 | P6988_RS00595 |
| *Vibrio coralliirubri* DSM 27495 | GCF_024347375.1 | OCV20_RS00330 | OCV20_RS00345 |
| *Vibrio cortegadensis* CECT 7227 | GCF_024347395.1 | OCV39_RS13735 | OCV39_RS13715 |
| *Vibrio crassostreae* LMG 22240 | GCF_024347415.1 | OC193_RS13615 | OC193_RS13600 |
| *Vibrio cyclitrophicus* ECSMB14105 | GCF_005144905.1 | FAZ90_RS02005 | FAZ90_RS02020 |

**Supplementary Table S1.** (Continued).

| **Strain** | **Assembly** | **16S rDNA locus** | **23S rDNA locus** |
| --- | --- | --- | --- |
| *Vibrio diabolicus* NV27 | GCF_020717785.1 | LJY22_RS13940 | LJY22_RS13920 |
| *Vibrio diazotrophicus* ATCC 33466 | GCF_038452265.1 | AAGA51_RS15230 | AAGA51_RS15205 |
| *Vibrio echinoideorum* DSM 107264 | GCF_024347455.1 | OCV36_RS00660 | OCV36_RS00675 |
| *Vibrio europaeus* NPI-1 | GCF_013154935.1 | HOO69_RS00025 | HOO69_RS00040 |
| *Vibrio fluvialis* ATCC 33809 | GCF_001558415.2 | AL536_RS44120 | AL536_RS44135 |
| *Vibrio fortis* LMG 21557 | GCF_024347475.1 | OCV50_RS01520 | OCV50_RS01535 |
| *Vibrio furnissii* FDAARGOS_777 | GCF_006364355.1 | FIU11_RS03140 | FIU11_RS03160 |
| *Vibrio gallaecicus* CECT 7244 | GCF_024347495.1 | OCU78_RS01610 | OCU78_RS01615 |
| *Vibrio gallicus* CIP 107863 | GCF_024346875.1 | OCU28_RS11210 | OCU28_RS11205 |
| *Vibrio gangliei* DSM 104291 | GCF_026001925.1 | Vgang_RS11520 | Vgang_RS11510 |
| *Vibrio gazogenes* PB1 | GCF_002196515.1 | BSQ33_RS11080 | BSQ33_RS11100 |
| *Vibrio gigantis* LMG 22741 | GCF_022371215.1 | MID13_RS00190 | MID13_RS00215 |
| *Vibrio harveyi* ATCC 33843 | GCF_000770115.1 | LA59_RS03605 | LA59_RS03620 |
| *Vibrio hyugaensis* 090810a | GCF_002906655.1 | C1S74_RS11855 | C1S74_RS11880 |
| *Vibrio inusitatus* LMG 23434 | GCF_024346935.1 | OCU48_RS00105 | OCU48_RS00125 |
| *Vibrio ishigakensis* C1 | GCF_024347675.1 | Pcarn_RS13095 | Pcarn_RS13080 |
| *Vibrio japonicus* JCM 31412 | GCF_024582835.1 | NP165_RS02140 | NP165_RS02150 |
| *Vibrio jasicida* 090810c | GCF_002887615.1 | C1S73_RS12600 | C1S73_RS12615 |
| *Vibrio kanaloae* R17 | GCF_001995825.2 | BTD91_RS06720 | BTD91_RS06725 |
| *Vibrio lentus* LMG 21034 | GCF_024347555.1 | OCV49_RS00105 | OCV49_RS00125 |
| *Vibrio mangrovi* CECT 7927 | GCF_024346955.1 | OCU74_RS01965 | OCU74_RS01980 |
| *Vibrio maritimus* BH16 | GCF_021441885.1 | LY387_RS13400 | LY387_RS13385 |
| *Vibrio mediterranei* QT6D1 | GCF_002214345.1 | BSZ05_RS07110 | BSZ05_RS07130 |
| *Vibrio metoecus* 08-2459 | GCF_009665275.1 | EWA65_RS01705 | EWA65_RS01715 |
| *Vibrio metschnikovii* 9502-00 | GCF_009763765.1 | GPX86_RS06415 | GPX86_RS06400 |
| *Vibrio mimicus* SCCF01 | GCF_001767355.1 | vm_RS00245 | vm_RS00260 |
| *Vibrio natriegens* CCUG 16374 | GCF_001680085.1 | BA894_RS14650 | BA894_RS14635 |

**Supplementary Table S1.** (Continued).

| **Strain** | **Assembly** | **16S rDNA locus** | **23S rDNA locus** |
| --- | --- | --- | --- |
| *Vibrio navarrensis* 2462-79 | GCF_009763725.1 | GPY30_RS15930 | GPY30_RS15915 |
| *Vibrio neonatus* JCM 21521 | GCF_024346975.1 | OCU38_RS12150 | OCU38_RS12135 |
| *Vibrio nigripulchritudo* SFn1 | GCF_000801275.2 | VIBNI_RS15940 | VIBNI_RS15915 |
| *Vibrio ostreae* OG9-811 | GCF_019226825.1 | KNV97_RS06340 | KNV97_RS06350 |
| *Vibrio owensii* SH14 | GCF_001310575.2 | APZ19_RS14270 | APZ19_RS14255 |
| *Vibrio palustris* CECT 9027 | GCF_024346995.1 | OCU30_RS00290 | OCU30_RS00300 |
| *Vibrio panuliri* JCM 19500^T^ | GCF_009938205.1 | GZK95_RS14305 | GZK95_RS14290 |
| *Vibrio paracholerae* NCTC 30 | GCF_900538065.1 | D5R51_RS01645 | D5R51_RS01660 |
| *Vibrio parahaemolyticus* RIMD 2210633 | GCF_000196095.1 | VP_RS14515 | VP_RS14510 |
| *Vibrio pectenicida* LMG 19642 | GCF_024347015.1 | OCV27_RS14190 | OCV27_RS14170 |
| *Vibrio penaeicida* IFO 15640 | GCF_019977755.1 | LDO37_RS17905 | LDO37_RS17885 |
| *Vibrio plantisponsor* CECT 7581 | GCF_024347035.1 | OCV04_RS01145 | OCV04_RS01155 |
| *Vibrio pomeroyi* LMG 20537 | GCF_024347595.1 | OCV12_RS01350 | OCV12_RS01355 |
| *Vibrio ponticus* DSM 16217 | GCF_009938225.1 | GZN30_RS10035 | GZN30_RS10020 |
| *Vibrio porteresiae* MSSRF30 | GCF_024347055.1 | OCV11_RS15855 | OCV11_RS15850 |
| *Vibrio rarus* LMG 23674 | GCF_024347075.1 | OCU56_RS11455 | OCU56_RS11450 |
| *Vibrio rhizosphaerae* LMG 23790 | GCF_024347095.1 | OCV37_RS00270 | OCV37_RS00280 |
| *Vibrio rotiferianus* B64D1 | GCF_002214395.1 | BSZ04_RS24475 | BSZ04_RS24495 |
| *Vibrio ruber* LMG 23124 | GCF_024347115.1 | OCU42_RS00315 | OCU42_RS00325 |
| *Vibrio rumoiensis* FERM P-14531 | GCF_002218045.2 | VRUMOI_RS09815 | VRUMOI_RS09790 |
| *Vibrio scophthalmi* VS-12 | GCF_001685465.1 | VSVS12_RS00280 | VSVS12_RS00265 |
| *Vibrio sinaloensis* YA2 | GCF_023195835.1 | MTO69_RS13475 | MTO69_RS13450 |
| *Vibrio spartinae* CECT 9026 | GCF_024347135.1 | OCU60_RS02290 | OCU60_RS02310 |
| *Vibrio splendidus* 2_C04b | GCF_003345295.1 | DUN60_RS14540 | DUN60_RS14565 |
| *Vibrio syngnathi* K08M4 | GCF_002119525.1 | K08M4_RS00710 | K08M4_RS00715 |
| *Vibrio taketomensis* C4III291 | GCF_009938185.1 | GZI12_RS01000 | GZI12_RS01005 |

**Supplementary Table S1.** (Continued).

| **Strain** | **Assembly** | **16S rDNA locus** | **23S rDNA locus** |
| --- | --- | --- | --- |
| *Vibrio tapetis* subsp. *tapetis* CECT4600^T^ | GCF_900233005.1 | VTAP4600_RS00060 | VTAP4600_RS00075 |
| *Vibrio tarriae* 2521-89 | GCF_002216685.1 | CEQ48_RS19535 | CEQ48_RS19560 |
| *Vibrio tasmaniensis* LMG 20012^T^ | GCF_024347635.1 | OCV44_RS00675 | OCV44_RS00690 |
| *Vibrio toranzoniae* CECT 7225^T^ | GCF_024347655.1 | OCU50_RS13650 | OCU50_RS13635 |
| *Vibrio tritonius* JCM 16457 | GCF_038695265.1 | AAD080_RS00255 | AAD080_RS00275 |
| *Vibrio tubiashii* ATCC 19109^T^ | GCF_000772105.1 | IX91_RS14485 | IX91_RS14495 |
| *Vibrio vulnificus* FORC_037 | GCF_002204915.1 | FORC37_RS00300 | FORC37_RS00315 |
| *Vibrio zhugei* HBUAS61001 | GCF_003716875.1 | EAE30_RS16575 | EAE30_RS16560 |
| *Vibrio ziniensis* ZWAL4003 | GCF_011064285.1 | G5S32_RS02010 | G5S32_RS02025 |
| *Escherichia coli* O157:H7 str. Sakai | GCF_000008865.2 | ECs_R0014 | ECs_R0015 |
| *Salmonella bongori* NCTC 12419 | GCF_000252995.1 | SBG_RS17885 | SBG_RS17895 |
| *Photobacterium* damselae subsp. *damselae* WMD-P2 | GCF_038086725.1 | WAX86_RS00040 | WAX86_RS00035 |

**Supplementary Table S2. *Vibrio* loci analyzed in this study.**

| **Locus** | **Description** | **Reference length (nt)*** |
| --- | --- | --- |
| Long universal single-copy genes | | |
| *alaS* | Alanyl-tRNA synthetase | 2583 |
| *aspS* | Aspartyl-tRNA synthetase | 1779 |
| *glnS* | Glutaminyl-tRNA synthetase or glutamine--tRNA ligase | 1671 |
| *gyrB* | DNA gyrase subunit B | 2418 |
| *leuS* | Leucyl-tRNA synthetase | 2574 |
| *metG* | Methionyl-tRNA synthetase | 2067 |
| *mfd* | Transcription-repair coupling factor (superfamily II helicase) | 3462 |
| *nrdA* | Ribonucleotide reductase alpha subunit | 2283 |
| *parC* | DNA topoisomerase IV subunit A | 2286 |
| *parE* | DNA topoisomerase IV subunit B | 1881 |
| *pnp* | Polyribonucleotide nucleotidyltransferase (polynucleotide phosphorylase) | 2136 |
| *polA* | DNA polymerase I, 3'-5' exonuclease and polymerase domains | 2796 |
| *recN* | DNA repair ATPase RecN | 1665 |
| *rpoB* | DNA-directed RNA polymerase, beta subunit/140 kD subunit | 4029 |
| *typA* | Predicted membrane GTPase TypA/BipA involved in stress response | 1830 |
| *uvrC* | Excinuclease UvrABC, nuclease subunit | 1833 |
| Synteny groups and subgroup in *Vibrio* | | |
| Group A | 5’-*secE*-*nusG*-*rplK*-*rplA*-*rplJ*-*rplL*-*rpoB*-3’ | 7729 |
| Group A.1 | 5’-*secE*-*nusG*-*rplK*-*rplA*-*rplJ*-*rplL*-3’ | 3450 |
| Group B | 5’-*lepA*-*lepB***-*rnc*-*era*-3’ | 4450 |
| Group C | 5’-*rpsB*-*tsf*-*pyrH*-*frr*-3’ | 3240 |
| Group D | 5’-*ruvA*-*ruvB*-3’ | 1641 |
| Group E | 5’-*rpsM*-*rpsK*-*rpsD*-*rpoA*-*rplQ*-3’ | 2837 |

* In *Vibrio parahaemolyticus* RIMD 2210633 (GCF_000196095.1).

** Gene *lepB* has not been proposed as a universal-single copy gene.

**Supplementary Table S3. Discarded loci due to lack of convergence during phylogenetic inference, or one or more quality criteria not being met.**

| **Locus** | **Description** | **Reference length (nt)*** |
| --- | --- | --- |
| Long universal single-copy genes | | |
| *argS* | Arginyl-tRNA synthetase | 1734 |
| *dnaG* | DNA primase (bacterial type) | 1761 |
| *dnaX* | DNA polymerase III, gamma/tau subunits | 2136 |
| *gyrA* | DNA topoisomerase (ATP-hydrolyzing) subunit A | 2637 |
| *infB* | Translation initiation factor IF-2, a GTPase | 2718 |
| *lepA* | Translation elongation factor EF-4, membrane-bound GTPase | 1794 |
| *pheT* | Phenylalanyl-tRNA synthetase beta subunit | 2406 |
| *recG* | RecG-like helicase | 2082 |
| *secA* | Preprotein translocase subunit SecA (ATPase, RNA helicase) | 2727 |
| Synteny groups in *Vibrio* | | |
| Group F | 5’-*rimM*-*trmD*-*rplS*-3’ | 1715 |
| Group G | 5’-*leuS*-*lptE***-*holA*-*rsfS*-3’ | 4721 |
| Group H | 5’-*nusA*-*infB*-*rbfA*-*truB*-3’ | 5583 |
| Group I | 5’-*rpsL*-*rpsG*-3’ | 947 |
| Group J | 5’-*dnaX*-*YbaB/EbfC*-*recR*-3’ | 3177 |
| Group K | 5’-*parE*-*parC*-3’ | 4170 |
| Group L | 5’-*pheS*-*pheT*-3’ | 3408 |
| Group M | 5’-*infC*-*rpmI***-*rplT*-3’ | 1260 |
| Group N | 5’-*tig*-*clpP***-*clpX*-3’ | 3389 |
| Group O | 3'- *ftsY*-5’\|5’-*rsmD*-3' | 2005 |
| Group P | 5’-*rpsJ*-*rplC*-*rplD*-*rplW*-*rplB*-*rpsS*-*rplV*-*rpsC*-*rplP*-*rpmC*-*rpsQ*-3’ | 4942 |
| Group Q | 5’-*rplN*-*rplX*-*rplE*-*rpsN*-*rpsH*-*rplF*-*rplR*-*rpsE*-3’ | 3438 |
| Group R | 5’-*rplO*-*secY*-3’ | 1790 |

* In *Vibrio parahaemolyticus* RIMD 2210633 (GCF_000196095.1).

** Genes not proposed as universal single-copy genes.

.

**Supplementary Table S4. Universal primers used to delimit 16S rDNA variable regions.**

| **Primer name** | **Usage in this study** | **Primer references** |
| --- | --- | --- |
| 27F | 5’ boundary of V1 | (Jian et al., 2019; Lane, 1991) |
| V2f | 5’ boundary of V2 | (Will et al., 2010) |
| 336R | 3’ boundary of V2 | (Weidner et al., 1996) |
| 337F | 5’ boundary of V3 | (Gwak and Rho, 2020) |
| 515F | 5’ boundary of V4 | (Caporaso et al., 2011) |
| 518R | 3’ boundary of V3 | (Muyzer et al., 1993) |
| 785F | 5’ boundary of V5 | (Kraneveld et al., 2012) |
| 806R | 3’ boundary of V4 | (Caporaso et al., 2011) |
| 907R | 3’ boundary of V5 | (Mao et al., 2012) |
| 1100F | 5’ boundary of V7 | (Liu et al., 2007) |
| 1100R | 3’ boundary of V6 | (Qing et al., 2024) |
| 1175R | 3’ boundary of V7  5’ boundary of V8 | (Kraneveld et al., 2012) |
| 1392wR | 3’ boundary of V8  5’ boundary of V9 | (Hoedt et al., 2024) |
| 1492R | 3’ boundary of V9 | (Jian et al., 2019; Lane, 1991) |

Abbreviations. V: Variable region.

**Supplementary Table S5. Partial 16S rDNA sequences analyzed in this study.**

| **Partial 16S rDNA sequence** | **Reference coordinates*** | **Reference length (nt)*** | **Boundary primers** |
| --- | --- | --- | --- |
| 16S rDNA V1-V2+V4-V9 | join(8..368,525..1521) | 1358 | 27F-336R + 515F-1492R |
| 16S rDNA V1-V2+V7-V9 | join(8..368,1110..1521) | 773 | 27F-336R + 1100F-1492R |
| 16S rDNA V1-V3+V5-V9 | join(8..544,795..1521) | 1264 | 27F-518R + 785F-1492R |
| 16S rDNA V1-V5 | (8..936) | 929 | 27F-907R |
| 16S rDNA V1-V6 | (8..1124) | 1117 | 27F-1100R |
| 16S rDNA V1-V6+V8-V9 | join(8..1124,1186..1521) | 1453 | 27F-1100R + 1175R-1492R |
| 16S rDNA V1-V7 | (8..1205) | 1187 | 27F-1175R |
| 16S rDNA V1-V7+V9 | join(8..1205,1403..1521) | 1317 | 27F-1175R + 1392wR-1492R |
| 16S rDNA V1-V8 | (8..1417) | 1410 | 27F-1392wR |
| 16S rDNA V2-V9 | (111..1521) | 1411 | V2f-1492R |
| 16S rDNA V3-V4 | (351..816) | 466 | 337F-806R |
| 16S rDNA V3-V6 | (351..1124) | 774 | 337F-1100R |
| 16S rDNA V3-V7 | (351..1205) | 855 | 337F-1175R |
| 16S rDNA V3-V9 | (351..1521) | 1171 | 337F-1492R |
| 16S rDNA V4-V9 | (525..1521) | 997 | 515F-1492R |

* Reference: *Vibrio parahaemolyticus* RIMD 2210633 (GCF_000196095.1), 16S rDNA locus VP_RS14515.

**Supplementary Table S6. Bayesian phylogenetic inference configuration for each alignment.**

| **Alignment** | **DNA substitution model** | **T** | **Number of generations** | **Sample frequency** |
| --- | --- | --- | --- | --- |
| 16S rDNA | HKY+I+G4  (Hasegawa et al., 1985) | 0.075 | 1 x 10^8^ | 4 x 10^4^ |
| 16S rDNA V1-V2+V4-V9 | SYM+I+G4  (Zharkikh, 1994) | 0.075 | 1 x 10^8^ | 4 x 10^4^ |
| 16S rDNA V1-V2+V7-V9 | SYM+I+G4 | 0.075 | 5 x 10^7^ | 2 x 10^4^ |
| 16S rDNA V1-V3+V5-V9 | GTR+I+G4  (Tavaré, 1986) | 0.075 | 5 x 10^7^ | 2 x 10^4^ |
| 16S rDNA V1-V5 | GTR+I+G4 | 0.050 | 5 x 10^7^ | 2 x 10^4^ |
| 16S rDNA V1-V6 | GTR+I+G4 | 0.075 | 5 x 10^7^ | 2 x 10^4^ |
| 16S rDNA V1-V6+V8-V9 | HKY+I+G4 | 0.100 | 5 x 10^7^ | 2 x 10^4^ |
| 16S rDNA V1-V7 | GTR+I+G4 | 0.075 | 5 x 10^7^ | 2 x 10^4^ |
| 16S rDNA V1-V7+V9 | HKY+I+G4 | 0.075 | 1 x 10^8^ | 4 x 10^4^ |
| 16S rDNA V1-V8 | SYM+I+G4 | 0.075 | 5 x 10^7^ | 2 x 10^4^ |
| 16S rDNA V2-V9 | HKY+I+G4 | 0.100 | 1 x 10^8^ | 4 x 10^4^ |
| 16S rDNA V3-V4 | GTR+I+G4 | 0.075 | 5 x 10^7^ | 2 x 10^4^ |
| 16S rDNA V3-V6 | GTR+I+G4 | 0.075 | 1 x 10^8^ | 4 x 10^4^ |
| 16S rDNA V3-V7 | HKY+I+G4 | 0.075 | 5 x 10^7^ | 2 x 10^4^ |
| 16S rDNA V3-V9 | HKY+I+G4 | 0.075 | 1 x 10^8^ | 4 x 10^4^ |
| 16S rDNA V4-V9 | HKY+I+G4 | 0.075 | 5 x 10^7^ | 2 x 10^4^ |
| 23S rDNA | GTR+I+G4 | 0.100 | 5 x 10^7^ | 2 x 10^4^ |
| *alaS* | GTR+I+G4 | 0.100 | 5 x 10^7^ | 2 x 10^4^ |
| *aspS* | GTR+I+G4 | 0.050 | 5 x 10^7^ | 2 x 10^4^ |
| *glnS* | GTR+I+G4 | 0.100 | 5 x 10^7^ | 2 x 10^4^ |
| *gyrB* | GTR+I+G4 | 0.100 | 5 x 10^7^ | 2 x 10^4^ |
| *leuS* | GTR+I+G4 | 0.075 | 5 x 10^7^ | 2 x 10^4^ |
| *metG* | GTR+I+G4 | 0.100 | 5 x 10^7^ | 2 x 10^4^ |
| *mfd* | GTR+I+G4 | 0.050 | 1 x 10^8^ | 4 x 10^4^ |
| *nrdA* | GTR+I+G4 | 0.050 | 5 x 10^7^ | 2 x 10^4^ |
| *parC* | GTR+I+G4 | 0.075 | 5 x 10^7^ | 2 x 10^4^ |
| *parE* | GTR+I+G4 | 0.100 | 5 x 10^7^ | 2 x 10^4^ |

**Supplementary Table S6.** (Continued).

| **Alignment** | **DNA substitution model** | **T** | **Number of generations** | **Sample frequency** |
| --- | --- | --- | --- | --- |
| *pnp* | GTR+I+G4 | 0.100 | 5 x 10^7^ | 2 x 10^4^ |
| *polA* | GTR+I+G4 | 0.075 | 5 x 10^7^ | 2 x 10^4^ |
| *recN* | GTR+I+G4 | 0.100 | 5 x 10^7^ | 2 x 10^4^ |
| *rpoB* | GTR+I+G4 | 0.100 | 1 x 10^8^ | 4 x 10^4^ |
| *typA* | GTR+I+G4 | 0.100 | 5 x 10^7^ | 2 x 10^4^ |
| *uvrC* | GTR+I+G4 | 0.100 | 5 x 10^7^ | 2 x 10^4^ |
| Group A | GTR+I+G4 | 0.100 | 1 x 10^8^ | 4 x 10^4^ |
| Group A.1 | GTR+I+G4 | 0.100 | 5 x 10^7^ | 2 x 10^4^ |
| Group B | GTR+I+G4 | 0.050 | 5 x 10^7^ | 2 x 10^4^ |
| Group C | GTR+I+G4 | 0.100 | 1 x 10^8^ | 4 x 10^4^ |
| Group D | GTR+I+G4 | 0.100 | 5 x 10^7^ | 2 x 10^4^ |
| Group E | GTR+I+G4 | 0.075 | 5 x 10^7^ | 2 x 10^4^ |

Abbreviations. HKY+I+G4: Hasegawa-Kishino-Yano model with a proportion of invariable sites and gamma distributed rate variation among sites. SYM+I+G4: Symmetric model with a proportion of invariable sites and gamma distributed rate variation among sites. GTR+I+G4: General time reversible model with a proportion of invariable sites and gamma distributed rate variation among sites. T: Temperature constant.

**Supplementary Table S7. Characteristics of USCGs included in the loci under study.** Functional categories and pathways were defined according to the information present in the NCBI COG database on September 12, 2024. The sixteen long USCGs analyzed as individual loci are indicated by an asterisk (*).

| **USCG** | **Gene product** | **Functional category** | **Pathway** |
| --- | --- | --- | --- |
| *alaS** | Alanyl-tRNA synthetase | Translation, ribosomal structure and biogenesis (J) | Aminoacyl-tRNA synthetases |
| *aspS** | Aspartyl-tRNA synthetase | Translation, ribosomal structure and biogenesis (J) | Aminoacyl-tRNA synthetases |
| *era* | GTPase Era, involved in 16S rRNA processing | Transcription (K) | N/A |
| *frr* | Ribosome recycling factor | Translation, ribosomal structure and biogenesis (J) | N/A |
| *glnS** | Glutaminyl-tRNA synthetase or glutamine--tRNA ligase | Translation, ribosomal structure and biogenesis (J) | Heme biosynthesis |
| *gyrB** | DNA gyrase subunit B | Replication, recombination and repair (L) | N/A |
| *lepA* | Translation elongation factor EF-4, membrane-bound GTPase | Translation, ribosomal structure and biogenesis (J) | N/A |
| *leuS** | Leucyl-tRNA synthetase | Translation, ribosomal structure and biogenesis (J) | Aminoacyl-tRNA synthetases |
| *metG** | Methionyl-tRNA synthetase | Translation, ribosomal structure and biogenesis (J) | Aminoacyl-tRNA synthetases |
| *mfd** | Transcription-repair coupling factor (superfamily II helicase) | Replication, recombination and repair (L), Transcription (K) | N/A |
| *nrdA** | Ribonucleotide reductase alpha subunit | Nucleotide transport and metabolism (F) | Pyrimidine salvage |
| *nusG* | Transcription termination / antitermination protein NusG | Transcription (K) | N/A |

**Supplementary Table S7.** (Continued).

| **USCG** | **Gene product** | **Functional category** | **Pathway** |
| --- | --- | --- | --- |
| *parC** | DNA topoisomerase IV subunit A | Replication, recombination and repair (L) | N/A |
| *parE** | DNA topoisomerase IV subunit B | Replication, recombination and repair (L) | N/A |
| *pnp** | Polyribonucleotide nucleotidyltransferase (polynucleotide phosphorylase) | Translation, ribosomal structure and biogenesis (J) | N/A |
| *polA** | DNA polymerase I, 3'-5' exonuclease and polymerase domains | Replication, recombination and repair (L) | N/A |
| *pyrH* | Uridylate kinase | Nucleotide transport and metabolism (F) | Pyrimidine biosynthesis |
| *recN** | DNA repair ATPase RecN | Replication, recombination and repair (L) | N/A |
| *rnc* | dsRNA-specific ribonuclease | Transcription (K) | N/A |
| *rplA* | Ribosomal protein L1 | Translation, ribosomal structure and biogenesis (J) | Ribosome 50S subunit |
| *rplJ* | Ribosomal protein L10 | Translation, ribosomal structure and biogenesis (J) | Ribosome 50S subunit |
| *rplK* | Ribosomal protein L11 | Translation, ribosomal structure and biogenesis (J) | Ribosome 50S subunit |
| *rplL* | Ribosomal protein L7/L12 | Translation, ribosomal structure and biogenesis (J) | Ribosome 50S subunit |
| *rplQ* | Ribosomal protein L17 | Translation, ribosomal structure and biogenesis (J) | Ribosome 50S subunit |

**Supplementary Table S7.** (Continued).

| **USCG** | **Gene product** | **Functional category** | **Pathway** |
| --- | --- | --- | --- |
| *rpoA* | DNA-directed RNA polymerase, alpha subunit/40 kD subunit | Transcription (K) | RNA polymerase |
| *rpoB** | DNA-directed RNA polymerase, beta subunit/140 kD subunit | Transcription (K) | RNA polymerase |
| *rpsB* | Ribosomal protein S2 | Translation, ribosomal structure and biogenesis (J) | Ribosome 30S subunit |
| *rpsD* | Ribosomal protein S4 or related protein | Translation, ribosomal structure and biogenesis (J) | Ribosome 30S subunit |
| *rpsK* | Ribosomal protein S11 | Translation, ribosomal structure and biogenesis (J) | Ribosome 30S subunit |
| *rpsM* | Ribosomal protein S13 | Translation, ribosomal structure and biogenesis (J) | Ribosome 30S subunit |
| *ruvA* | Holliday junction resolvasome RuvABC DNA-binding subunit | Replication, recombination and repair (L) | N/A |
| *ruvB* | Holliday junction resolvasome RuvABC, ATP-dependent DNA helicase subunit RuvB | Replication, recombination and repair (L) | N/A |
| *secE* | Preprotein translocase subunit SecE | Intracellular trafficking, secretion, and vesicular transport (U) | Sec pathway |
| *tsf* | Translation elongation factor EF-Ts | Translation, ribosomal structure and biogenesis (J) | Translation factors |
| *typA** | Predicted membrane GTPase TypA/BipA involved in stress response | Signal transduction mechanisms (T) | N/A |
| *uvrC** | Excinuclease UvrABC, nuclease subunit | Replication, recombination and repair (L) | N/A |

**Supplementary Table S8. Alignment length and number of variable sites of the multiple sequence alignments used for Bayesian phylogenetic inference.**

| **Alignment** | **Alignment length (nt)** | **Number of variable sites** |
| --- | --- | --- |
| 16S rDNA | 1592 | 319 |
| 16S rDNA V1-V2+V4-V9 | 1397 | 279 |
| 16S rDNA V1-V2+V7-V9 | 811 | 192 |
| 16S rDNA V1-V3+V5-V9 | 1303 | 271 |
| 16S rDNA V1-V5 | 965 | 203 |
| 16S rDNA V1-V6 | 1153 | 230 |
| 16S rDNA V1-V6+V8-V9 | 1492 | 296 |
| 16S rDNA V1-V7 | 1234 | 249 |
| 16S rDNA V1-V7+V9 | 1355 | 276 |
| 16S rDNA V1-V8 | 1447 | 287 |
| 16S rDNA V2-V9 | 1444 | 278 |
| 16S rDNA V3-V4 | 466 | 79 |
| 16S rDNA V3-V6 | 755 | 123 |
| 16S rDNA V3-V7 | 856 | 142 |
| 16S rDNA V3-V9 | 1175 | 207 |
| 16S rDNA V4-V9 | 1001 | 172 |
| 23S rDNA | 2925 | 624 |
| *alaS* | 2679 | 1628 |
| *aspS* | 1803 | 1009 |
| *glnS* | 1675 | 939 |
| *gyrB* | 2443 | 1329 |
| *leuS* | 2109 | 1183 |
| *metG* | 3622 | 2305 |
| *mfd* | 2295 | 1259 |
| *nrdA* | 2443 | 1329 |
| *parC* | 2295 | 1305 |
| *parE* | 1902 | 1047 |
| *pnp* | 2178 | 1145 |
| *polA* | 2901 | 1826 |

**Supplementary Table S8.** (Continued).

| **Alignment** | **Alignment length (nt)** | **Number of variable sites** |
| --- | --- | --- |
| *recN* | 1678 | 1174 |
| *rpoB* | 4035 | 1823 |
| *typA* | 1842 | 997 |
| *uvrC* | 1911 | 1322 |
| Group A | 8341 | 3980 |
| Group A.1 | 3919 | 1964 |
| Group B | 5403 | 2983 |
| Group C | 3862 | 2081 |
| Group D | 2021 | 1159 |
| Group E | 2884 | 1065 |

**Supplementary Table S9. Bayesian phylogenetic inference diagnostic metrics for each alignment.**

| **Alignment** | **Swap frequency between adjacent chains (%)** | **ASDSF** | **Average PSRF** |
| --- | --- | --- | --- |
| 16S rDNA | 19 - 35 | 0.0182 | 1.000 |
| 16S rDNA  V1-V2+V4-V9 | 17 - 33 | 0.0178 | 1.000 |
| 16S rDNA  V1-V2+V7-V9 | 14 - 32 | 0.0221 | 1.000 |
| 16S rDNA  V1-V3+V5-V9 | 19 - 35 | 0.0167 | 1.000 |
| 16S rDNA  V1-V5 | 35 - 49 | 0.0098 | 1.000 |
| 16S rDNA  V1-V6 | 18 - 34 | 0.0266 | 1.000 |
| 16S rDNA  V1-V6+V8-V9 | 11 - 25 | 0.0222 | 1.000 |
| 16S rDNA  V1-V7 | 16 - 32 | 0.0152 | 1.000 |
| 16S rDNA  V1-V7+V9 | 19 - 33 | 0.0182 | 1.001 |
| 16S rDNA  V1-V8 | 18 - 34 | 0.0124 | 1.000 |
| 16S rDNA  V2-V9 | 11 - 24 | 0.0125 | 1.000 |
| 16S rDNA  V3-V4 | 30 - 47 | 0.0113 | 1.000 |
| 16S rDNA  V3-V6 | 15 - 33 | 0.0185 | 1.000 |
| 16S rDNA  V3-V7 | 14 - 32 | 0.0194 | 1.001 |
| 16S rDNA  V3-V9 | 15 - 33 | 0.0154 | 1.000 |
| 16S rDNA  V4-V9 | 12 - 31 | 0.0127 | 1.000 |
| 23S rDNA | 18 - 28 | 0.0186 | 1.000 |
| *alaS* | 27 - 34 | 0.0078 | 1.000 |
| *aspS* | 56 - 60 | 0.0143 | 1.000 |
| *glnS* | 27 - 34 | 0.0095 | 1.000 |

**Supplementary Table S9.** (Continued).

| **Alignment** | **Swap frequency between adjacent chains (%)** | **ASDSF** | **Average PSRF** |
| --- | --- | --- | --- |
| *gyrB* | 26 - 35 | 0.0101 | 1.000 |
| *leuS* | 40 - 45 | 0.0042 | 1.000 |
| *metG* | 25 - 34 | 0.0166 | 1.000 |
| *mfd* | 55 - 60 | 0.0046 | 1.000 |
| *nrdA* | 53 - 60 | 0.0127 | 1.000 |
| *parC* | 39 - 44 | 0.0059 | 1.000 |
| *parE* | 26 - 35 | 0.0065 | 1.000 |
| *pnp* | 23 - 32 | 0.0102 | 1.000 |
| *polA* | 36 - 44 | 0.0089 | 1.000 |
| *recN* | 22 - 32 | 0.0059 | 1.000 |
| *rpoB* | 25 - 31 | 0.0125 | 1.000 |
| *typA* | 26 - 34 | 0.0222 | 1.000 |
| *uvrC* | 24 - 34 | 0.0095 | 1.000 |
| Group A | 26 - 33 | 0.0063 | 1.000 |
| Group A.1 | 15 - 33 | 0.0205 | 1.000 |
| Group B | 50 - 57 | 0.0290 | 1.001 |
| Group C | 22 - 34 | 0.0249 | 1.001 |
| Group D | 23 - 35 | 0.0095 | 1.000 |
| Group E | 35 - 43 | 0.0069 | 1.000 |

Abbreviations. ASDSF: Average standard deviation of split frequencies. PSRF: Potential scale reduction factor.

**Supplementary Table S10. Position in ordination space of phylogenetic trees from USCG loci and rDNA representative copies.** Coordinates represented as mean ± standard deviation. Data represented in Fig. 1.

| **Alignment** | **nRF distance** | | **wRF distance** | |
| --- | --- | --- | --- | --- |
|  | **NMDS1** | **NMDS2** | **NMDS1** | **NMDS2** |
| 16S rDNA | -0.6391 ± 0.0782 | -0.1978 ± 0.0599 | -0.5251 ± 0.1670 | -0.0093 ± 0.1735 |
| 23S rDNA | 0.0874 ± 0.1275 | -0.1709 ± 0.2052 | -0.0289 ± 0.1567 | -0.4108 ± 0.1406 |
| *alaS* | 0.0564 ± 0.0414 | -0.0165 ± 0.0572 | 0.0012 ± 0.0455 | 0.0531 ± 0.0553 |
| *aspS* | 0.1068 ± 0.1037 | 0.0028 ± 0.1348 | 0.0079 ± 0.0904 | 0.0768 ± 0.1116 |
| *glnS* | 0.0274 ± 0.0195 | 0.0185 ± 0.0236 | 0.0319 ± 0.0336 | 0.0414 ± 0.0353 |
| *gyrB* | 0.1250 ± 0.0649 | -0.0244 ± 0.1074 | -0.0419 ± 0.0803 | 0.0378 ± 0.1014 |
| *leuS* | 0.0436 ± 0.1062 | 0.0285 ± 0.1542 | -0.0755 ± 0.0711 | 0.0430 ± 0.0856 |
| *metG* | 0.1275 ± 0.0773 | -0.0502 ± 0.1100 | -0.0281 ± 0.0780 | 0.0258 ± 0.0934 |
| *mfd* | 0.0111 ± 0.0201 | 0.0278 ± 0.0258 | 0.0294 ± 0.0194 | 0.0156 ± 0.0190 |
| *nrdA* | 0.0004 ± 0.0139 | 0.0232 ± 0.0158 | 0.0280 ± 0.0276 | 0.0332 ± 0.0291 |
| *parC* | 0.0279 ± 0.0216 | -0.0114 ± 0.0199 | -0.0099 ± 0.0208 | 0.0123 ± 0.0208 |
| *parE* | 0.0135 ± 0.0259 | 0.0015 ± 0.0230 | -0.0100 ± 0.0343 | 0.0305 ± 0.0407 |
| *pnp* | 0.1847 ± 0.0944 | -0.0343 ± 0.1274 | -0.0509 ± 0.1700 | 0.0311 ± 0.2410 |
| *polA* | -0.0110 ± 0.0138 | 0.0253 ± 0.0148 | 0.0153 ± 0.0188 | 0.0120 ± 0.0193 |
| *recN* | 0.0103 ± 0.0188 | 0.0251 ± 0.0225 | 0.0722 ± 0.0438 | -0.0021 ± 0.0462 |
| *rpoB* | 0.0034 ± 0.0211 | 0.0057 ± 0.0249 | 0.1123 ± 0.0674 | 0.0126 ± 0.0676 |
| *typA* | -0.3272 ± 0.0568 | 0.2949 ± 0.0635 | -0.1348 ± 0.1534 | 0.0356 ± 0.2035 |
| *uvrC* | 0.0152 ± 0.0286 | 0.0384 ± 0.0320 | 0.0345 ± 0.0295 | 0.0063 ± 0.0324 |
| Group A | 0.0190 ± 0.0152 | 0.0030 ± 0.0187 | 0.1189 ± 0.0638 | -0.0082 ± 0.0596 |
| Group A.1 | 0.0220 ± 0.0194 | 0.0051 ± 0.0206 | 0.1521 ± 0.0879 | -0.0280 ± 0.0865 |
| Group B | 0.0291 ± 0.0122 | 0.0043 ± 0.0187 | 0.0449 ± 0.0222 | 0.0137 ± 0.0217 |
| Group C | 0.0488 ± 0.0342 | -0.0319 ± 0.0377 | 0.0621 ± 0.0542 | -0.0305 ± 0.0661 |
| Group D | 0.0175 ± 0.0290 | 0.0209 ± 0.0339 | 0.0260 ± 0.0241 | 0.0302 ± 0.0262 |
| Group E | 0.0003 ± 0.0347 | 0.0126 ± 0.0372 | 0.1684 ± 0.1162 | -0.0219 ± 0.1220 |

Abbreviations. nRF: Normalized Robinson-Foulds. wRF: Weighted Robinson-Foulds. NMDS1: First dimension of non-metric multidimensional scaling ordination. NMDS2: Second dimension of non-metric multidimensional scaling ordination.

**Supplementary Table S11. Position in ordination space of phylogenetic trees from USCG loci, rDNA representative copies, and partial 16S rDNA sequences without a single variable region.** Coordinates represented as mean ± standard deviation. Data represented in Fig. 2.

| **Alignment** | **nRF distance** | | **wRF distance** | |
| --- | --- | --- | --- | --- |
|  | **NMDS1** | **NMDS2** | **NMDS1** | **NMDS2** |
| 16S rDNA | -0.3069 ± 0.0134 | 0.0154 ± 0.0153 | 0.2729 ± 0.0372 | 0.0000 ± 0.0423 |
| 16S rDNA  V1-V2+V4-V9 | -0.3057 ± 0.0165 | 0.0332 ± 0.0645 | 0.2957 ± 0.0379 | 0.0062 ± 0.1353 |
| 16S rDNA  V1-V3+V5-V9 | -0.3073 ± 0.0146 | 0.0135 ± 0.0137 | 0.2868 ± 0.0382 | 0.0037 ± 0.0427 |
| 16S rDNA  V1-V6+V8-V9 | -0.2822 ± 0.0167 | 0.0137 ± 0.0089 | 0.2604 ± 0.0367 | 0.0003 ± 0.0413 |
| 16S rDNA  V1-V7+V9 | -0.3161 ± 0.0130 | 0.0108 ± 0.0209 | 0.3212 ± 0.0378 | 0.0025 ± 0.0527 |
| 16S rDNA  V1-V8 | -0.3142 ± 0.0137 | 0.0160 ± 0.0151 | 0.2920 ± 0.0378 | 0.0027 ± 0.0425 |
| 16S rDNA  V2-V9 | -0.2828 ± 0.0156 | 0.0026 ± 0.0433 | 0.2553 ± 0.0349 | 0.0090 ± 0.0952 |
| 23S rDNA | 0.0803 ± 0.0232 | 0.0282 ± 0.1480 | -0.0424 ± 0.0463 | -0.2765 ± 0.0472 |
| *alaS* | 0.0925 ± 0.0091 | 0.0259 ± 0.0250 | -0.0791 ± 0.0220 | 0.0264 ± 0.0297 |
| *aspS* | 0.1285 ± 0.0254 | 0.0240 ± 0.0617 | -0.0983 ± 0.0343 | 0.0333 ± 0.0635 |
| *glnS* | 0.1053 ± 0.0054 | 0.0039 ± 0.0147 | -0.0999 ± 0.0180 | 0.0232 ± 0.0179 |
| *gyrB* | 0.1503 ± 0.0154 | 0.0132 ± 0.0387 | -0.0760 ± 0.0282 | 0.0203 ± 0.0683 |
| *leuS* | 0.1109 ± 0.0247 | 0.0088 ± 0.0794 | -0.0355 ± 0.0218 | 0.0201 ± 0.0556 |
| *metG* | 0.1194 ± 0.0207 | 0.0421 ± 0.0494 | -0.0479 ± 0.0264 | 0.0143 ± 0.0487 |
| *mfd* | 0.1056 ± 0.0054 | -0.0069 ± 0.0146 | -0.0796 ± 0.0128 | 0.0102 ± 0.0124 |
| *nrdA* | 0.0743 ± 0.0071 | -0.0180 ± 0.0069 | -0.0818 ± 0.0175 | 0.0189 ± 0.0153 |
| *parC* | 0.0720 ± 0.0079 | 0.0107 ± 0.0083 | -0.0405 ± 0.0134 | 0.0075 ± 0.0087 |
| *parE* | 0.0732 ± 0.0093 | 0.0063 ± 0.0112 | -0.0575 ± 0.0176 | 0.0163 ± 0.0207 |
| *pnp* | 0.1653 ± 0.0203 | 0.0340 ± 0.0536 | -0.1246 ± 0.0426 | 0.0103 ± 0.1623 |
| *polA* | 0.0675 ± 0.0065 | -0.0176 ± 0.0068 | -0.0644 ± 0.0128 | 0.0082 ± 0.0114 |
| *recN* | 0.1058 ± 0.0060 | -0.0074 ± 0.0128 | -0.1088 ± 0.0174 | 0.0029 ± 0.0331 |
| *rpoB* | 0.0709 ± 0.0071 | -0.0037 ± 0.0117 | -0.1362 ± 0.0251 | 0.0109 ± 0.0368 |
| *typA* | -0.0492 ± 0.0178 | -0.2570 ± 0.0224 | 0.0218 ± 0.0266 | 0.0121 ± 0.1296 |

**Supplementary Table S11.** (Continued).

| **Alignment** | | **nRF distance** | | **wRF distance** | |
| --- | --- | --- | --- | --- | --- |
|  |  | **NMDS1** | **NMDS2** | **NMDS1** | **NMDS2** |
|  | *uvrC* | 0.1111 ± 0.0082 | -0.0089 ± 0.0186 | -0.0861 ± 0.0166 | 0.0070 ± 0.0202 |
|  | Group A | 0.0830 ± 0.0057 | 0.0012 ± 0.0095 | -0.1405 ± 0.0210 | 0.0019 ± 0.0374 |
|  | Group A.1 | 0.0905 ± 0.0090 | -0.0026 ± 0.0120 | -0.1645 ± 0.0285 | -0.0041 ± 0.0554 |
|  | Group B | 0.0912 ± 0.0044 | 0.0010 ± 0.0094 | -0.0890 ± 0.0150 | 0.0078 ± 0.0133 |
|  | Group C | 0.0945 ± 0.0097 | 0.0154 ± 0.0248 | -0.1078 ± 0.0252 | -0.0138 ± 0.0317 |
|  | Group D | 0.0993 ± 0.0094 | 0.0043 ± 0.0181 | -0.0846 ± 0.0156 | 0.0179 ± 0.0135 |
|  | Group E | 0.0729 ± 0.0113 | -0.0021 ± 0.0167 | -0.1612 ± 0.0395 | 0.0005 ± 0.0896 |

Abbreviations. nRF: Normalized Robinson-Foulds. wRF: Weighted Robinson-Foulds. NMDS1: First dimension of non-metric multidimensional scaling ordination. NMDS2: Second dimension of non-metric multidimensional scaling ordination.

**Supplementary Table S12. Position in ordination space of phylogenetic trees from USCG loci, rDNA representative copies, and different partial 16S rDNA sequences.** Coordinates represented as mean ± standard deviation. Data represented in Supplementary Fig. S2.

| **Alignment** | **nRF distance** | | **wRF distance** | |
| --- | --- | --- | --- | --- |
|  | **NMDS1** | **NMDS2** | **NMDS1** | **NMDS2** |
| 16S rDNA | -0.1736 ± 0.0214 | -0.0324 ± 0.0257 | -0.1375 ± 0.0308 | 0.0432 ± 0.0409 |
| 16S rDNA  V1-V2+V4-V9 | -0.1587 ± 0.0249 | -0.0765 ± 0.0579 | -0.1477 ± 0.0397 | 0.0868 ± 0.0940 |
| 16S rDNA  V1-V2+V7-V9 | -0.1653 ± 0.0483 | -0.1906 ± 0.1072 | -0.0672 ± 0.0566 | 0.2011 ± 0.1642 |
| 16S rDNA  V1-V5 | -0.2603 ± 0.0267 | -0.0708 ± 0.0843 | -0.1809 ± 0.0507 | 0.0894 ± 0.1441 |
| 16S rDNA  V1-V6 | -0.2252 ± 0.0210 | -0.0209 ± 0.0346 | -0.1879 ± 0.0397 | 0.0452 ± 0.0577 |
| 16S rDNA  V1-V7 | -0.2197 ± 0.0201 | -0.0199 ± 0.0334 | -0.1913 ± 0.0363 | 0.0452 ± 0.0501 |
| 16S rDNA  V3-V4 | -0.2224 ± 0.0553 | 0.1451 ± 0.1552 | -0.2736 ± 0.1025 | -0.1901 ± 0.1749 |
| 16S rDNA  V3-V6 | -0.2378 ± 0.0251 | 0.1019 ± 0.0519 | -0.2539 ± 0.0527 | -0.0766 ± 0.0907 |
| 16S rDNA  V3-V7 | -0.2439 ± 0.0213 | 0.0854 ± 0.0451 | -0.2445 ± 0.0451 | -0.0658 ± 0.0748 |
| 16S rDNA  V3-V9 | -0.2217 ± 0.0291 | 0.0496 ± 0.0513 | -0.2171 ± 0.0448 | -0.0176 ± 0.0696 |
| 16S rDNA  V4-V9 | -0.1699 ± 0.0668 | -0.0289 ± 0.2462 | -0.1606 ± 0.0780 | 0.0325 ± 0.2326 |
| 23S rDNA | 0.1032 ± 0.0187 | 0.0277 ± 0.0837 | 0.1257 ± 0.0473 | -0.0289 ± 0.1443 |
| *alaS* | 0.1012 ± 0.0084 | 0.0069 ± 0.0239 | 0.0771 ± 0.0164 | -0.0143 ± 0.0253 |
| *aspS* | 0.1107 ± 0.0157 | 0.0137 ± 0.0473 | 0.0802 ± 0.0238 | -0.0180 ± 0.0457 |
| *glnS* | 0.1062 ± 0.0076 | -0.0028 ± 0.0109 | 0.0936 ± 0.0166 | -0.0058 ± 0.0212 |
| *gyrB* | 0.1446 ± 0.0113 | 0.0043 ± 0.0239 | 0.0729 ± 0.0201 | -0.0049 ± 0.0391 |
| *leuS* | 0.1262 ± 0.0149 | -0.0018 ± 0.0443 | 0.0428 ± 0.0146 | -0.0069 ± 0.0261 |
| *metG* | 0.1141 ± 0.0156 | 0.0016 ± 0.0446 | 0.0540 ± 0.0179 | -0.0016 ± 0.0288 |
| *mfd* | 0.1144 ± 0.0075 | -0.0015 ± 0.0104 | 0.0815 ± 0.0120 | -0.0006 ± 0.0108 |
| *nrdA* | 0.0806 ± 0.0076 | -0.0024 ± 0.0112 | 0.0857 ± 0.0162 | -0.0034 ± 0.0173 |

**Supplementary Table S12.** (Continued).

| **Alignment** | | **nRF distance** | | **wRF distance** | |
| --- | --- | --- | --- | --- | --- |
|  |  | **NMDS1** | **NMDS2** | **NMDS1** | **NMDS2** |
|  | *parC* | 0.0844 ± 0.0070 | 0.0056 ± 0.0103 | 0.0539 ± 0.0104 | -0.0013 ± 0.0082 |
|  | *parE* | 0.0803 ± 0.0081 | -0.0018 ± 0.0109 | 0.0633 ± 0.0136 | -0.0019 ± 0.0159 |
|  | *pnp* | 0.1548 ± 0.0128 | 0.0065 ± 0.0327 | 0.1133 ± 0.0351 | -0.0221 ± 0.0866 |
|  | *polA* | 0.0687 ± 0.0060 | -0.0030 ± 0.0087 | 0.0695 ± 0.0118 | -0.0010 ± 0.0102 |
|  | *recN* | 0.1146 ± 0.0082 | -0.0002 ± 0.0106 | 0.1101 ± 0.0166 | -0.0048 ± 0.0170 |
|  | *rpoB* | 0.0776 ± 0.0061 | 0.0043 ± 0.0086 | 0.1196 ± 0.0216 | -0.0214 ± 0.0237 |
|  | *typA* | 0.0411 ± 0.0170 | 0.0012 ± 0.1426 | 0.0184 ± 0.0218 | -0.0015 ± 0.0597 |
|  | *uvrC* | 0.1149 ± 0.0086 | 0.0012 ± 0.0132 | 0.0878 ± 0.0153 | 0.0012 ± 0.0138 |
|  | Group A | 0.0892 ± 0.0065 | -0.0083 ± 0.0081 | 0.1303 ± 0.0203 | -0.0080 ± 0.0183 |
|  | Group A.1 | 0.0941 ± 0.0101 | -0.0051 ± 0.0089 | 0.1517 ± 0.0248 | -0.0080 ± 0.0237 |
|  | Group B | 0.0933 ± 0.0060 | -0.0002 ± 0.0072 | 0.0869 ± 0.0141 | -0.0073 ± 0.0116 |
|  | Group C | 0.1063 ± 0.0085 | 0.0024 ± 0.0196 | 0.1052 ± 0.0202 | -0.0114 ± 0.0209 |
|  | Group D | 0.0973 ± 0.0094 | 0.0109 ± 0.0125 | 0.0815 ± 0.0140 | -0.0064 ± 0.0161 |
|  | Group E | 0.0810 ± 0.0093 | -0.0012 ± 0.0135 | 0.1571 ± 0.0306 | -0.0151 ± 0.0382 |

Abbreviations. nRF: Normalized Robinson-Foulds. wRF: Weighted Robinson-Foulds. NMDS1: First dimension of non-metric multidimensional scaling ordination. NMDS2: Second dimension of non-metric multidimensional scaling ordination.

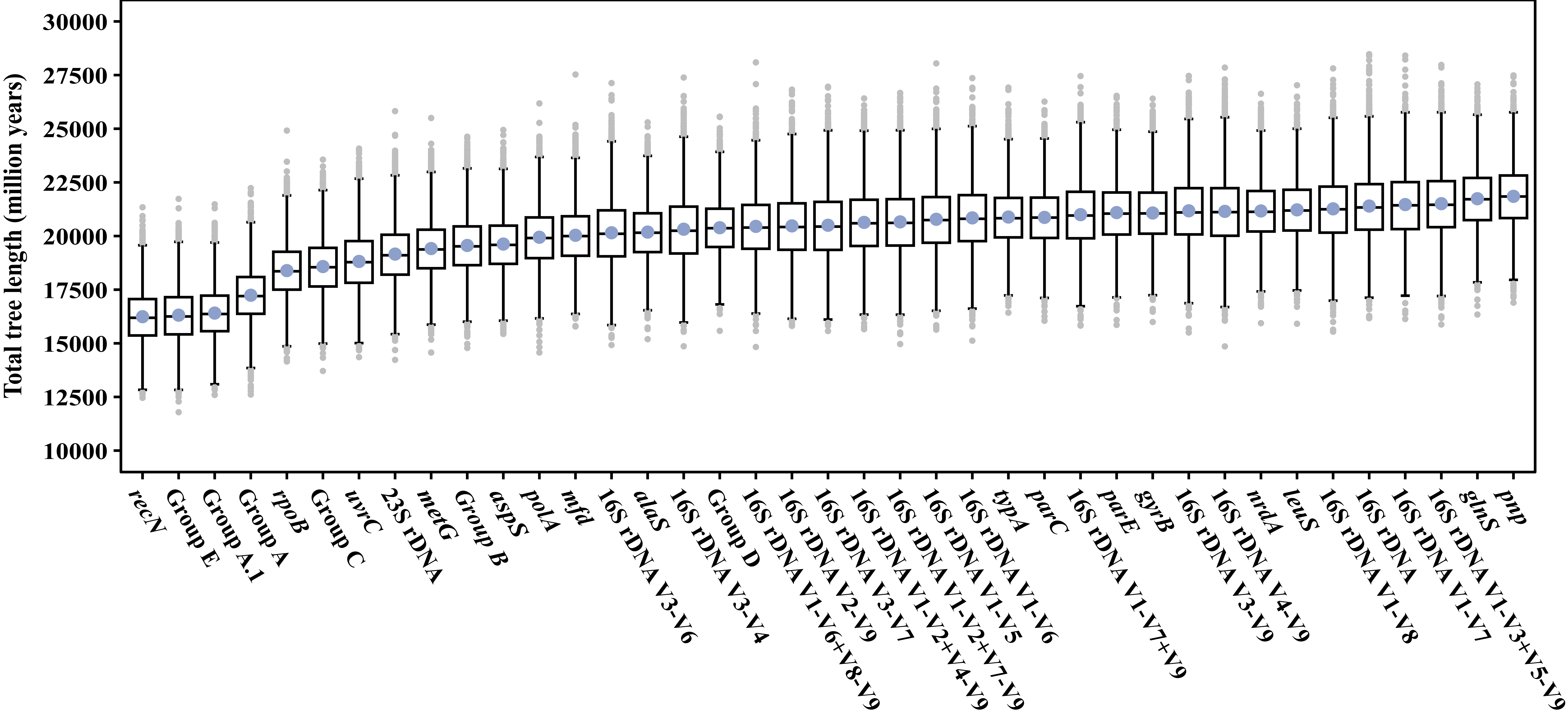

**Supplementary Fig. S1. Total tree length of the posterior distribution produced by each alignment.** Blue dots represent the mean total tree length of the posterior distribution of trees. Grey dots represent outliers.

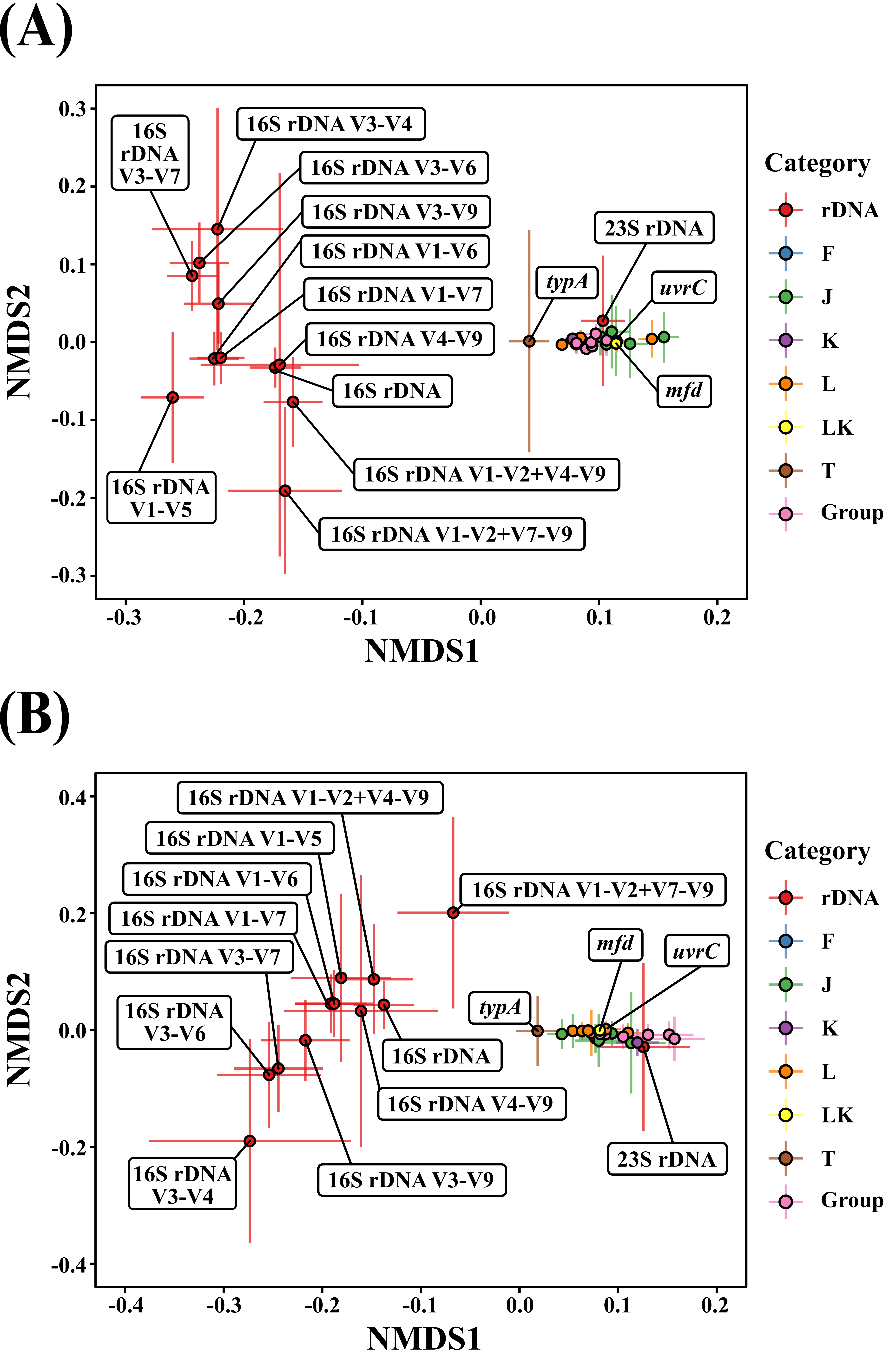

**Fig. S2. NMDS analysis of phylogenetic congruence between USCG loci, rDNA representative copies, and different partial 16S rDNA sequences.** Mean position of loci (dots) with standard deviation bars in two dimensions. **(A)** Ordination of distances among trees calculated according to the nRF metric, which is influenced by differences in tree topology. Stress (mean ± standard deviation): 0.12 ± 0.01. **(B)** Ordination of distances among trees calculated according to the wRF metric, which incorporates information on variation in tree branch lengths. Stress (mean ± standard deviation): 0.15 ± 0.02. Categories: rDNA represents 16S and 23S rDNA loci; F, J, K, L, LK and T represent individual USCGs according to their functional categories (Supplementary Table S7); and synteny groups as indicated.
